# Spatiotemporal dynamics of successive activations across the human brain during simple arithmetic processing

**DOI:** 10.1101/2023.11.22.568334

**Authors:** Pedro Pinheiro-Chagas, Clara Sava-Segal, Serdar Akkol, Amy Daitch, Josef Parvizi

## Abstract

Previous neuroimaging studies have offered unique insights about the spatial organization of activations and deactivations across the brain, however these were not powered to explore the exact timing of events at the subsecond scale combined with precise anatomical source information at the level of individual brains. As a result, we know little about the order of engagement across different brain regions during a given cognitive task. Using experimental arithmetic tasks as a prototype for human-unique symbolic processing, we recorded directly across 10,076 brain sites in 85 human subjects (52% female) using intracranial electroencephalography (iEEG). Our data revealed a remarkably distributed change of activity in almost half of the sampled sites. Notably, an orderly successive activation of a set of brain regions - anatomically consistent across subjects-was observed in individual brains. Furthermore, the temporal order of activations across these sites was replicable across subjects and trials. Moreover, the degree of functional connectivity between the sites decreased as a function of temporal distance between regions, suggesting that information is partially leaked or transformed along the processing chain. Furthermore, in each activated region, distinct neuronal populations with opposite activity patterns during target and control conditions were juxtaposed in an anatomically orderly manner. Our study complements the prior imaging studies by providing hitherto unknown information about the timing of events in the brain during arithmetic processing. Such findings can be a basis for developing mechanistic computational models of human-specific cognitive symbolic systems.

**Significance statement:** Our study elucidates the spatiotemporal dynamics and anatomical specificity of brain activations across >10,000 sites during arithmetic tasks, as captured by intracranial EEG. We discovered an orderly, successive activation of brain regions, consistent across individuals, and a decrease in functional connectivity as a function of temporal distance between regions. Our findings provide unprecedented insights into the sequence of cognitive processing and regional interactions, offering a novel perspective for enhancing computational models of cognitive symbolic systems.

## INTRODUCTION

Past studies in humans and non-human primates have provided important information about regional activations during arithmetic processing (1–8). More specifically, neuropsychological and neuroimaging studies have uncovered a static and coarse functional map of activations in the brain, which in the context of arithmetic processing, includes a network of regions within the posterior parietal cortex (PPC) - presumably engaged in numerosity representation and manipulation(9–14) - and the ventral temporal cortex (VTC) - presumably involved in the recognition of numerical symbols (15–19). The engaged PPC sites are further grouped into at least three functionally diverse sub-regions(20): the horizontal portion of the intraparietal sulcus (IPS), which is strongly engaged during number comparison(14) and calculation(21), the superior parietal lobe (SPL), which hosts a topographical representation of quantities(22) and is involved in visuospatial processing and orienting of attention during calculation(12, 23, 24), and the left angular gyrus (AG), which is supposedly involved in verbal number processing such as multiplication fact-retrieval(25, 26) (but see(27, 28)). Performing a simple arithmetic calculation would therefore depend on an interplay between different subregions of the PPC, between PPC and VTC, and between these regions and other auxiliary brain regions that are required for performing executive functions and working memory (e.g., dorsolateral prefrontal cortex, DLPFC), declarative and semantic memory formation (e.g., medial temporal lobe, MTL), and the allocation of attentional resources for goal-directed problem solving (e.g., various prefrontal cortical regions, PFC)(29).

While we have learned a great deal of information from the extant evidence, our knowledge has been primarily based on group-averaged correlational data that could not simultaneously resolve the spatial and temporal dynamics of the activity at the individual brain level. As a result, the precise anatomical location, temporal dynamics of activity and how these regions collaboratively work together to perform calculations are still poorly understood. This is partially due to the low temporal resolution and low signal-to-noise ratio of signals obtained with noninvasive neuroimaging methods. While the lower temporal resolution of fMRI precludes assessments of fast spatiotemporal dynamics across these sites, the lower signal-to-noise ratio of M/EEG often require a group-based analysis that blurs the anatomical precision of the data at the single-subject level.

In the past few years, studies using intracranial electroencephalography (iEEG) have provided several novel findings that are advancing our understanding of the spatiotemporal dynamics of processing in the human brain. While the iEEG method is not suited for fine-grained studies of neuronal activity at the local level, it is a remarkable tool for simultaneous recording of averaged neuronal population activities across remote regions of the brain(30). Using this method, our group has confirmed the existence of a specialized area in the posterior inferior temporal gyrus (pITG) in the human VTC that selectively responds to numerals as compared to other morphometrically and semantically similar symbols(17). Subsequent studies have shown that the pITG sites are functionally connected with neuronal populations around the intraparietal sulcus (IPS), found to be associated with magnitude representation and manipulation(31, 32) and modulated by arithmetic problem-size(13, 32). A further study demonstrated that responses in the pITG during calculation are format independent (i.e. equal responses for numerals and number words)(33). These results have suggested that the pITG may be involved in functions beyond digit recognition. Although our recent iEEG studies provided precise information about the anatomical location and temporal dynamics of activity between the VTC and IPS hubs of activity, they were only focused on a single pair of brain regions and relied on observations in a small number of subjects - due to the rarity of recordings in these areas of the brain.

In the current study, we leveraged the power of multisite iEEG recordings across a relatively large number of human subjects. Crucially, several of our subjects had simultaneous recordings across the key regions of the brain known to be involved during mathematical processing. This allowed us to investigate the spatiotemporal dynamics of arithmetic processing at the level of single subject and with single trial precision. We had several predictions based on previous studies. Based on neuropsychological data showing the heterogeneity of behavioral deficits following focal(34) and network brain lesions(35), we hypothesized the existence of juxtaposed neuronal populations with preferential responses for math and non-math tasks. Extending our prior work to a larger number of regions (31, 33, 36), we predicted that sites with a high degree of preferential response to math processing will show a particular signature of activity profile, which can be observed as a sharp increase in activation following the calculation stage of the task. Complementary, we expected to observe an antagonistic pattern of deactivation across regions of the default mode network at a similar processing stage, which has never been demonstrated. Lastly, expanding our previous findings with just a few electrodes (31), we predicted the existence of a canonical temporal order of activations along the math network, following regions in the ventral temporal, lateral parietal and dorsolateral prefrontal cortices. In addition to these specific hypotheses, our study was poised to gain novel insights into the precise anatomical organization within individual brains, the temporal dynamics, and functional connectivity during arithmetic calculations. Our findings have the potential to serve as a foundational platform for developing mechanistic computational models of uniquely human cognitive abilities, thus greatly advancing our understanding of mathematical cognition.

## MATERIALS AND METHODS

### Participants

We recorded electrocorticography data from 85 human subjects (52% females; see Extended Data Table 1 - Subject’s demographics, basic neuropsychological information and task completion) with medically refractory epilepsy who were implanted with intracranial electrodes as a part of their presurgical evaluation at Stanford University. Each subject provided informed consent to participate in the study, which was approved by the Stanford Institutional Review Board. Subjects were monitored for approximately 6-10 days following surgery, during which they participated in our study. Electrode location and number was determined by the neurosurgeons for clinical needs. Data from 16 subjects of the present cohort was already published elsewhere (17, 31–33, 37).with a center-to-center interelectrode spacing of 10 mm, or a 5 mm spacing for high-density arrays.

### Electrophysiology Recording

Each subject was implanted with grids and/or strips of subdural platinum electrodes (AdTech Medical Instruments Corporation), whose locations were determined purely for clinical reasons. For depth electrodes (stereo-EEG), the diameter was .86 mm, height was 2.29 mm and the distance between the centers of two adjacent electrodes was 5 mm. For subdural grids and strips (ECoG) electrodes, the diameter was usually 2.3 mm with inter-electrode spacing of 10mm, 7mm, or 5mm for higher density arrays.

### Data acquisition and analysis

Data were recorded with a multichannel recording system (Tucker Davis Technologies). Data were acquired with a band pass filter of 0.5-300 Hz and sampling rate of 1525.88 Hz. An electrode outside the seizure zone with the most silent electrocorticographic activity was selected as an online reference during acquisition. After data collection, we implemented a preprocessing pipeline to minimize noise in the electrophysiological signals. Initially, signals were downsampled to 1000 Hz. We then applied notch filters at 60, 120, and 180 Hz to the downsampled signals to remove electrical line noise. Subsequently, we identified and excluded noisy channels from further analyses. These were defined as channels with raw amplitude exceeding five times or falling below one-fifth of the median raw amplitude across all channels, or those exhibiting more than three times the median number of spikes across all channels (spikes were defined as jumps between consecutive data points larger than 80 µV). We also excluded electrodes whose overall power was five or more standard deviations above or below the mean power across channels, and those whose power spectrum deviated from the normal 1/f power spectrum, based on visual inspection. Electrodes marked by the clinical team as having epileptiform activity were also excluded from subsequent analyses, along with the noisy electrodes.” All non-excluded channels were then notch filtered at 60 Hz and harmonics to remove electric interference, then re-referenced to the mean of the filtered signals of the non-excluded channels. The re-referenced signal at each electrode was then band-pass filtered between 70 and 180 Hz (high frequency broadband, HFB) using sequential 10 Hz width band-pass windows (70-80 Hz, 80-90 Hz, etc.), using two-way, zero-lag, FIR filters. The instantaneous amplitude of each band-limited signal was computed by taking the modulus of the Hilbert Transformed signal. The amplitude of each 10 Hz band signal was normalized by its own mean, then these normalized amplitude time series were averaged together, yielding a single amplitude time-course for the HFB band. Data was analyzed using Matlab R2020_b with a custom-made pipeline: https://github.com/pinheirochagas/lbcn_preproc.

All analyses were conducted at the level of individual electrodes, and a subset (ROL, functional connectivity, and feature modulation) were done at the single-trial level. The group analyses were conducted at the second level, incorporating information computed at the trial and electrode levels. This approach allowed us to maintain the high resolution of our data while also enabling us to draw broader conclusions across subjects and regions.

### Electrode localization

We used iELVis toolbox in order to localize electrodes (38). All subjects had pre-implantation T1-weighted MRI and post-implantation CT scans following the electrode implantation. T1-weighted MRI scan was used to generate cortical surface and subcortical segmentation using recon-all command of Freesurfer v6.0.0 (39). The post-implant CT scan was aligned to the pre-implant MRI using the *flirt* from the Oxford Centre for Functional MRI of the Brain Software Library (40, 41) or using *bbregister* from Freesurfer (42) to get the best results. We then manually labeled each electrode on the T1-registered CT image using BioImage Suite (43). For subdural grid and strip electrodes (but not for depth electrodes), we used one of two brainshift correction methods in order to get the closest results to the intraoperative pictures (44, 45). To combine data across subjects, electrodes were meticulously determined by a highly trained neuroanatomist, J.P., who considered the anatomical landmarks and morphology of individual brains. Importantly, the neuroanatomist was completely blind to the results.

### Functional preferentiality

Our analyses were focused on task-induced changes in HFB activity (70-180 Hz), due to its high correlation with local spiking activity and the fMRI BOLD signal (46, 47). Additionally, the HFB signal is proven to have sufficient anatomical resolution to capture independent activity from each electrode, even when multiple electrodes are located within the same grid or brain region (47, 48). This is based on the understanding that the HFB signal reflects localized neural processes and can vary significantly across closely spaced electrodes. However, we acknowledge that this is a complex issue and that some degree of spatial autocorrelation may exist in the data due to shared inputs or local network effects (49).

We classified all sites based on their relative responses during target (arithmetic) vs. control (memory) trials. *Math involved* sites were defined as those with significantly different HFB activity during arithmetic trials compared to the pre-stimulus baseline period (see below). If the trial HFB activity was higher than baseline, we considered the site *math-active* and if the trial HFB activity was lower than baseline, we considered the site *math deactivated. Math preferential* sites were defined as those that satisfied three criteria: 1) significantly higher trial HFB compared to baseline; 2) significantly greater trial HFB activity during math compared to the non-math condition (autobiographical memory/language); 3) no significant difference in trial HFB activity during the non-math condition as compared to baseline. Math and non-math active sites were defined as those with significantly higher trial HFB compared to baseline in both math and non-math conditions and with no significant difference in trial HFB activity between math and non-math conditions. *Non-math involved*, *non-math active*, *non-math deactivated* and *non-math preferential* were classified using equivalent criteria. The trial and baseline periods for the comparisons used in the Simultaneous Calculation task were 100 - 1,000 ms following the stimulus presentation and - 200 – 0ms before stimulus presentation, respectively. The trial window was selected not only because all necessary information for performing the calculation (in the math condition) or retrieving the memory (in the non-math condition) is presented simultaneously, but also because this 1-second window is where the majority of the signal was concentrated. For the Sequential Calculation task, the trial and baseline periods were 2,100 to 4,300 ms following the first stimulus and -500 – 0ms before the first stimulus, respectively.

On the other hand, the Sequential Calculation task has a different structure with information presented in a staggered manner. Consequently, the trial window for this task is longer (2,200 ms) and begins 100 ms after the presentation of the third stimulus. This timing allows subjects to gather all necessary information before starting the calculation or memory retrieval process. The varying baseline durations were chosen based on the intertrial interval, which was 200 ms for the Simultaneous task and 500 for the Sequential task. This procedure ensured that the baseline period did not overlap with any task stimuli. Note that this is not circular to the response profile findings **(Figures 2 and 3)**, since the analysis comparing the HFB within different stages of calculation was performed post-hoc to the functional preferentiality analysis. Unpaired permutation tests were run to test for differences in HFB power between different task conditions, while paired permutation tests were run to test for a difference in HFB power between a task condition and baseline. All p-values in all analyses were FDR corrected by the total number of sites within each subject, using the procedure introduced by (50).

**Figure 1.**
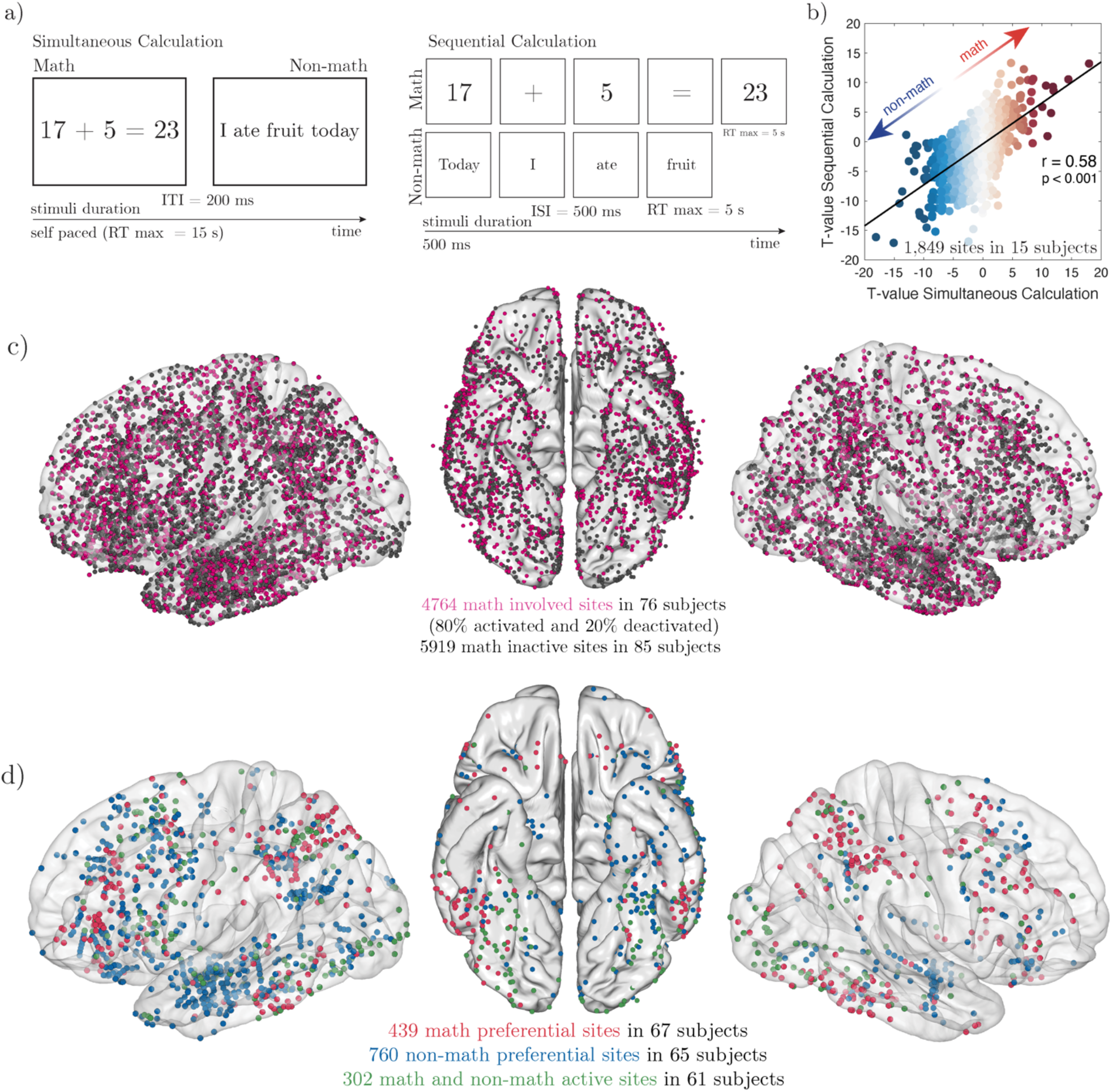
A) Design of the Simultaneous and Sequential Calculation tasks. B) Correlation of the T-value from the comparison between math vs. non-math averaged activity in each site from all the 15 subjects who completed both tasks. C) Electrode coverage combining all 10,076 recoding sites across 85 subjects plotted on a common brain (fsaverage). Each black dot represents one electrode. Most electrodes (62%) were ECoG and located on the left hemisphere (59%). D) Selectivity map for math and non-math (autobiographical memory/language) across all electrodes during the Simultaneous and Sequential Calculation tasks combined. Electrodes are colored by selectivity: math preferential in red, non-math preferential in blue and sites equally activated for both man and non-math in green. See Figures 1-(1–8).

**Figure 2.**
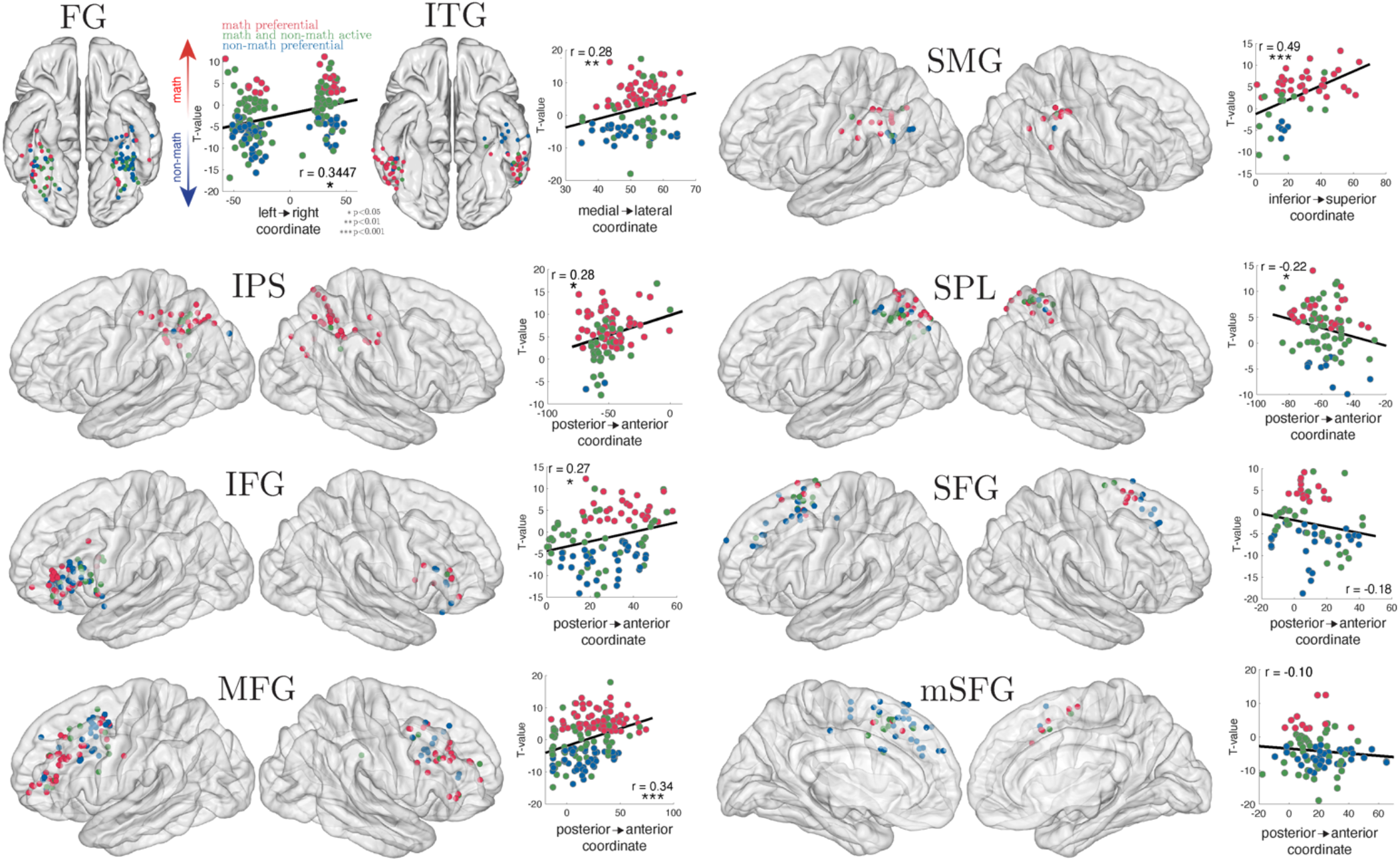
Anatomical localization of the preferential sites in the main nine hubs of the math network, combining the Simultaneous and Sequential Calculation tasks. While A, B, and C show electrodes in the standard MNI brains, scatter plots are from data in the individual anatomical space. The scatterplots show the correlation between the t-scores comparing math vs. non-math and the particular coordinate within each hub that best segregates between math and non-math preference in *the native individual anatomical space*. The correlation significance was FDR corrected for multiple comparisons, considering the 36 comparisons (9 regions x 4 coordinate axes: left/right, medial/lateral, posterior/anterior and inferior/superior). This anatomical relationship may not be as clear once individual subject’s data are transferred to standard MNI space. Fusiform gyrus, FG; inferior temporal gyrus, ITG; supramarginal gyrus, SMG; inferior parietal sulcus, IPS; superior parietal lobe, SPL; inferior frontal gyrus, IFG; superior frontal gyrus, SFG; middle frontal gyrus, MFG; medial superior frontal gyrus, mSFG. See Figure 1-4, Figure 1-5, Figure 1-6.

**Figure 3.**
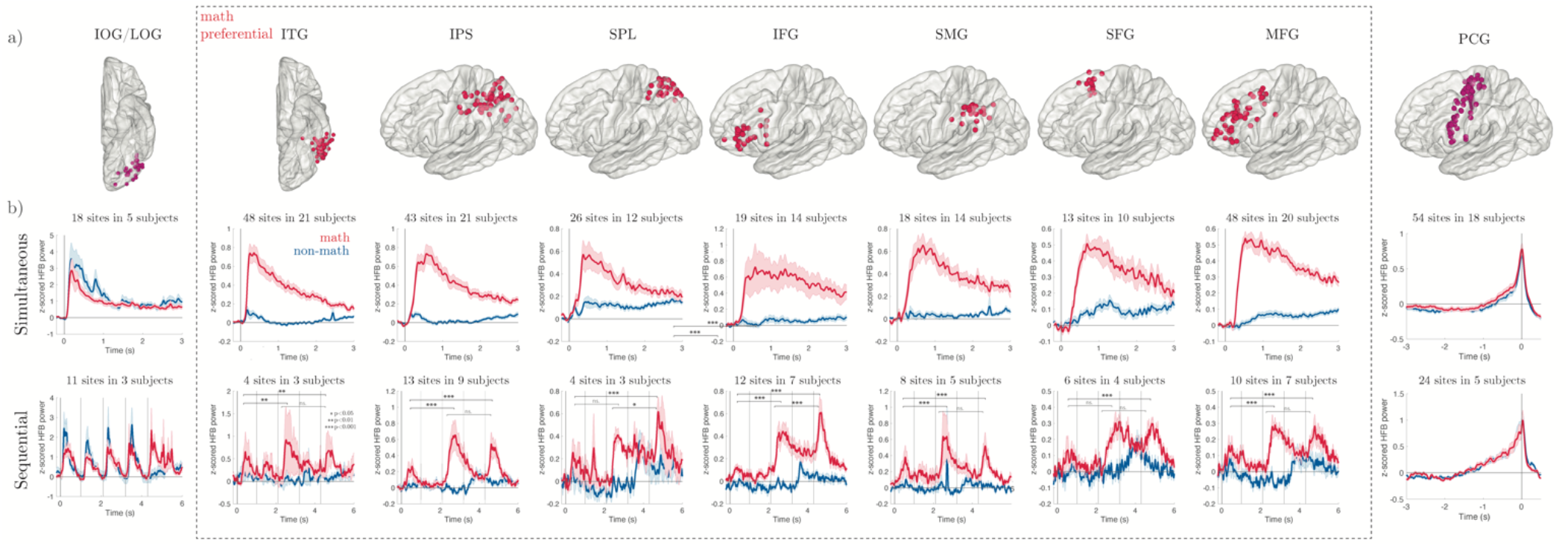
Profile of electrophysiological activity across the arithmetic network. A) Electrode localization of the main 7 regions that show strongest math selectivity in both Simultaneous and Sequential Calculation tasks (inside the dotted line), as well as 2 control regions that show no math selectivity (generic visual activations in the lateral occipital cortex and motor activations in the precentral gyrus). Shaded error bars represent the standard error of the mean across electrodes. B) HFB time courses for the Simultaneous and Sequential calculation tasks. Inferior occipital gyrus, IOG; lateral occipital gyrus, LOG; precentral gyrus, PCG. See Videos 1 and 2.

### Response onset latency analysis (ROL)

Based on our previously published methods (51), we implemented a method to estimate the onset of the task-induced trial-by-trial HFB response at each site. We first normalized the signal to the peak amplitude and applied a sliding window with 30 ms bins with 28 ms overlap. We then calculated the signal average and standard deviation in a baseline time window of [−200 0] ms before onset across trials in each condition. We then identified 25 consecutive bins in which the average HFB power z-score was one standard deviation above the baseline average. The earliest time point of the first bin in this sequence was marked as the onset for a specific trial. All analyses were done on a trial-by-trial basis. Sites in which the ROL could not be calculated for more than 50% of the trials were discarded from the analysis.

### Cross-correlation

For each pair of electrodes, we ran a cross-correlation analysis (function xcorr in Matlab) with max lag = [-2, 2] seconds from stimulus onset in the Simultaneous Calculation task. Next, we randomly shuffled trial order and re-calculated the cross correlation. We repeated this process 2,000 times and subtracted the averaged permuted shuffled trial order from the averaged *Pearson’*s r trace from preserved trial order. If two regions are functionally connected, the difference between ordered minus shuffled trials (*Δr_peak_*) should be high and positive. And if one region A is leading and another region B is lagging, the lag of the peak (*Δr_lag_)* should be positive.

### Simultaneous Calculation Task

Subjects were presented with visual stimuli that were either math statements (arithmetic calculations: i.e., “48 + 5 = 57”) or non-math statements (autobiographical memory: i.e., “I ate fruit yesterday”). Subjects were instructed to make true/false judgements about each statement or equation by pressing two keypad buttons. Each arithmetic statement (addition) was always composed of a two-digit operand (ranging from 10 to 87), a single-digit operand (ranging from 1 to 9), and a two-digit proposed result. In half of the trials, the proposed result was correct. The task was self-paced, but subjects had maximum of 15 s to make these judgements. A 200 ms intertrial period separated the trials. The trials were interspersed with fixation periods (5 sec), during which subjects were simply asked to fixate at a crosshair in the center of the screen. Subjects completed around 2 blocks. Each block had 80 trials (40 math and 40 non-math, randomly sampled from the stimuli list), which lasted ∼15 minutes. Stimuli were not repeated across blocks. Within each block, math and non-math statements were presented in a randomized order. (See Extended Data Table 2 - Stimuli list for the Simultaneous Calculation task for the full list of stimuli).

### Sequential Calculation Task

Subjects were presented with a series of visual stimuli that were either math statements or non-math statements, like the ones presented in the Simultaneous Calculation Task. Subjects were instructed to make true/false judgements about each statement or equation by pressing two keypad buttons. Each stimulus within a statement was presented one numeral/symbol/word at a time (i.e., “1” “+” “2” “=” “3”; “I” “ate” “fruit” “yesterday”). Half of the math trials were presented as math symbols (“1+2=3”) and half using words (“one plus two equals three”). In half of the math trials, the proposed result was correct. Each stimulus was presented for 500 ms with a 500 ms interstimulus period. After the last stimulus was presented, subjects had 5 s to respond. A 500 ms of intertrial period separated the trials. Subjects completed between 2-4 blocks. Each block consisted of 48 trials (24 math and 24 non-math, randomly sampled from the stimuli list), lasting ∼10 minutes. Stimuli were not repeated across blocks. Within each block, math and non-math statements were presented in a randomized order. See Extended Data Table 2 - Stimuli list for the Simultaneous Calculation task for the full list of stimuli.

We chose the distinction math vs. non-math, because for this paper we were not particularly focused on the precise cognitive functions underlying the non-math condition beyond that their lack of involvement in math; a similar terminology has been adopted in our recent studies. The non-math statements essentially represent a reading task with autobiographical memory content (19, 52).

## RESULTS

### Subjects and Recording Electrodes

We recruited subjects with focal epilepsy implanted with intracranial electrodes as part of their presurgical evaluation. Electrodes were implanted in these subjects solely for clinical reasons and subjects were monitored in private hospital rooms for 1-2 weeks. Our research studies were performed while subjects were comfortably resting in their hospital bed during their clinical monitoring. In total, 85 human subjects, including 49 cases with subdural grids (ECoG), 42 with depth electrodes (sEEG), and 6 cases with both sEEG and ECoG. All subjects were implanted with electrodes made by *AdTech*. A total of 10,076 recording sites were included: 56% in the left hemisphere and 56% ECoG) throughout the last 12 years. Subjects performed simple arithmetic sums presented in two different methods: In the first experiment, also referred to as Task I or “*Simultaneous Calculation*”, subjects judged the accuracy of full math statements (arithmetic equations: e.g., “16 + 8 = 22”) or full non-math statements (autobiographical memory: e.g., “Today, I had a beer”) that appeared in randomized order on a computer screen. In the second task, i.e., Task II, or “*Sequential Calculation*”, subjects viewed similar statements but with items presented serially, or one at a time (e.g., “Today”, “I” “had” “a beer” or “16”, “+”, “8”, “=”, “22”) (**Figure 1A**).

### Behavioral Performance

Subjects demonstrated high performance in the math and non-math conditions for both the simultaneous and sequential tasks. We classified trials as incorrect when subjects got the answer wrong, or when they had extreme RTs (<200MS or >10s for the Simultaneous and >5s for the Sequential math condition), since that suggested either a lack of attention or engagement. For the Simultaneous math condition, subjects accurately responded to 84.9% of the trials (std = 13.11%), with a mean RT = 3,723 ms, std = 1,283 ms (RT calculated from onset of full statement). For the Sequential math condition, subjects accurately responded to 83.8% of the trials (std = 8.4%), with a mean RT = 1,332 ms, std = 495 ms (RT calculated from the onset of the last item, i.e. “22” in “16+8=22”). In the non-math condition, since these involved subjective questions, accuracy could not be assessed. However, the RT for the Simultaneous non-math condition was mean = 2,635, std = 956, and for the Sequential non-math condition it was mean = 1,652, std = 471. Statistical analysis revealed significant RT differences between the math and non-math conditions across both tasks. In Simultaneous tasks, responses were slower in math condition by 1,088 ms, t(68) = 5.65, p < 0.0001, In contrast, subjects were faster for the math condition in the Sequential tasks by 320 ms, t(53) = 3.44, p < 0.001. Incorrect trials were excluded from all further analyses.

To model RTs, we calculated stepwise regression models for each participant with several arithmetic features that are classically associated with problem difficulty as predictors. Take a proposed problem like “7+35=44”, the *min operand* is the smallest operand (“7”), the *max operand* is the largest operand (“35”), the *cross-decade* is true, since the correct result “42” lies on the decade “40” that is different from the one of the max operand which is “30”; the *magnitude order* is either smaller operand + larger operand or larger operand + smaller operand (which was only possible to model in the Simultaneous Calculation task, because all stimuli of the Sequential Calculation task were larger operand + smaller operand); the *format* is a digit (in the Sequential Calculation task it can also be number words, such as “seven plus thirty five equals forty four”) and the *absolute deviant* is the modulus of the difference between the correct (“42”) and proposed (“44”) results, which in this case is “2”. In the Simultaneous Calculation task, results revealed that the cross-decade variable was overall the best predictor of RT (significant in 46 subjects, 70%), followed by min operand (significant in 36 subjects, 55%), absolute deviant (significant in 28 subjects, 42%), max operand (significant in 11 subjects, 17%) and magnitude order (significant in 9 subjects, 14%). In the Sequential Calculation task, we found that the best predictor of RT was the min operand (significant in 17 subjects, 65%), followed by format (significant in 15 subjects, 58%), absolute deviant (significant in 12 subjects, 46%), cross-decade (significant in 5 subjects, 19%), max operand (significant in 3 subjects, 12%).

These behavioral results corroborate previous findings using a variety of paradigms, such as verification, production, and number-to-position, in which the *min* operand and cross-decade are consistently found to be good predictors of RT (53–56), thus providing convergent evidence that these arithmetic features are robust and reproducible indices of problem difficulty.

### Functional Anatomical Organization

To investigate which sites were engaged during the math condition, we calculated band-limited power of signals recorded the High Frequency Broadband (HFB) (70-180Hz, normalized to the pre-stimulus baseline) as a reliable measure of “regional engagement” - as reviewed elsewhere (30).

By comparing the HFB activity during the calculation stage against the pre-stimulus baseline period in all 10,076 recording sites across 85 subjects, we identified that 4,764 sites (45%) were significantly involved during arithmetic calculations (**Figure 1C**). These sites were distributed throughout the entire mantle of the cerebral cortex in both hemispheres. Out of these involved sites, 3,811 (80%) were significantly activated (i.e., post-stimuli activity increased as compared to the baseline period, see Methods) and 953 (20%) were significantly deactivated (i.e., post-stimuli activity decreased as compared to the baseline period), (see **Figure 4**).

**Figure 4.**
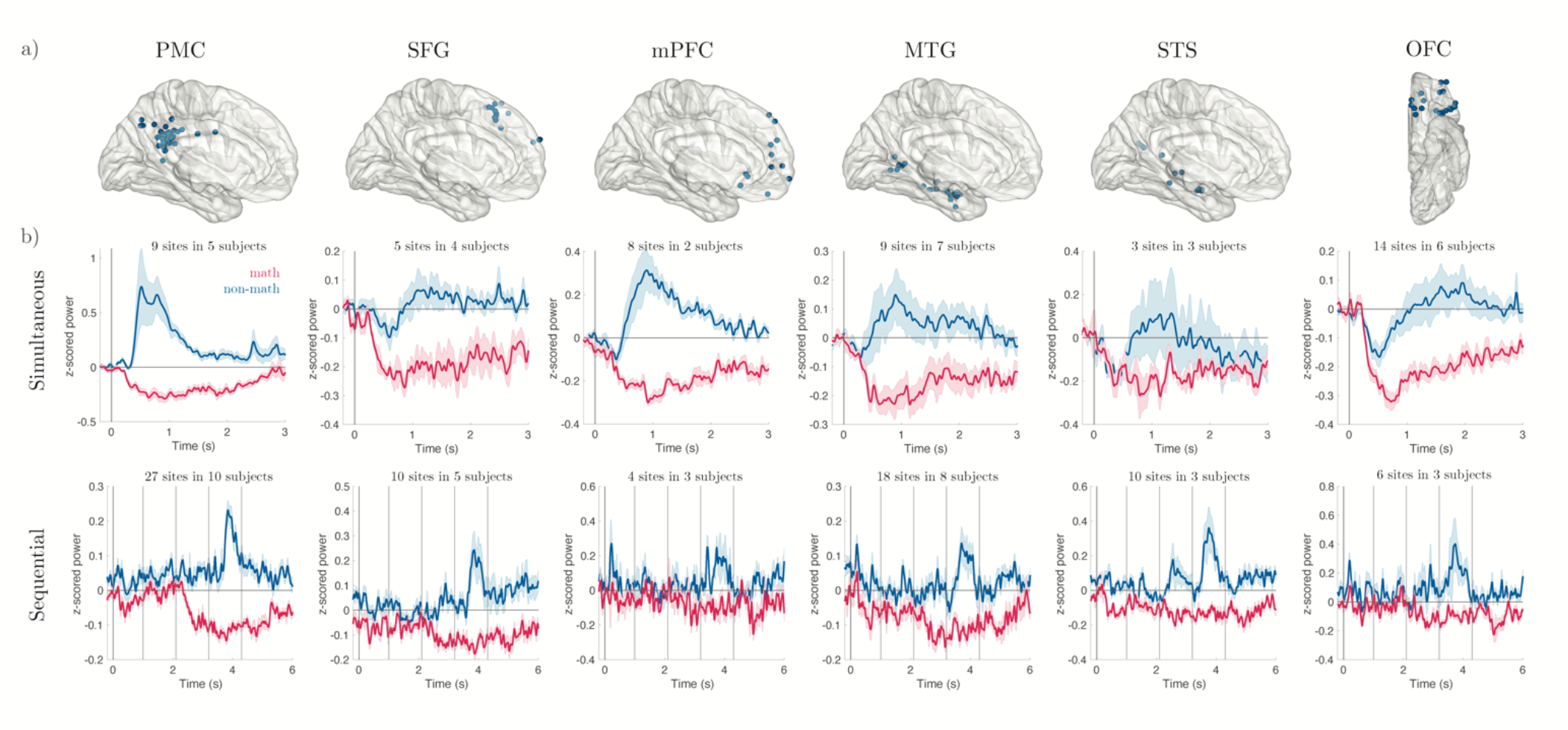
Profile of electrophysiological activity of region that were deactivated during math. A) Electrode localization of the main 6 regions that show strongest math decreased activity in both Simultaneous and Sequential Calculation tasks. B) HFB time courses for the Simultaneous and Sequential calculation tasks. Shaded error bars represent the standard error of the mean across electrodes.

As noted, the main goal of the current study was to study the dynamics of engagement and interaction across neuronal populations located in different regions of the brain during arithmetic processing. To address this aim, we focused our analysis on a selected set of brain regions that displayed the highest degree of activations during the arithmetic trials. In line with neuroimaging studies, we identified sites in each subject that exhibited significant activations (i.e., HFB responses) compared to baseline and higher activations during the math condition compared to non-math condition. It is important to note that this does not imply that the selected sites were engaged strictly in arithmetic computation *per se*. Instead, this subtractive approach (activity during math compared to activity during non-math control condition) was used to make our analyses similar to the ones employed in the majority of neuroimaging studies and to identify brain regions where peaks of activations are found.

Across all subjects, we identified 439 sites (9.21%, in 67 subjects) with preferential activity during math as compared to the non-math condition (**Figure 1D**). The t-scores from the comparison of induced HFB power during math compared to non-math conditions were highly correlated when calculated from Task I and Task II (i.e., simultaneous vs. sequential calculation, r = 0.55, p < 0.001, df=1,848, n = 15 subjects, 1,849 recording sites) (**Figure 1B**). This clearly suggests that the preferential responses in these sites were driven by the task content (i.e., arithmetic calculation) rather than by the task structure or demand.

Math preferential sites could be dissociated from the set of anatomical regions with preferential responses during non-math (N=760 in 65 subjects). The anatomical location of the sites with preferential activation during arithmetic processing were remarkably consistent across task format (i.e., in both Simultaneous and Sequential tasks) and across individual brains. We selected nine regions of interest for a deeper analysis based on the following criteria: 1. regions that have been implicated in mathematical processing in previous fMRI and neuropsychological studies; 2. regions that contained a substantial number of electrodes (i.e., n>10), prioritizing the Simultaneous Calculation task, given the significant coverage and emphasis on this task in our study. We acknowledge that this selection process may seem arbitrary, and we understand that despite the extensive nature of our database, it is not feasible to investigate all brain regions with the same level of precision. The selected ROIs were: fusiform gyrus (FG), inferior temporal gyrus (ITG), intraparietal sulcus (IPS), superior parietal lobule (SPL), supramarginal gyrus, inferior (IFG), middle (MFG) and superior frontal gyrus (SFG), and medial superior frontal (mSFG) gyrus (**Figure 2**). In each of these broad regions we could find both math and non-math preferential neuronal populations. However, consistent anatomical relationships could be observed between math and non-math activated sites. To confirm that, we plotted the T-value of the comparison between math vs. non-math and the electrode’s coordinate in the native individual anatomical space, using the coordinate axis that best captured this relationship. We found that functional preference was organized in these regions. In the FG, math responsive sites were located primarily in the right hemisphere while non-math activated sites were located predominantly in the left hemisphere. Moreover, the math activated sites were located more lateral to non-math sites in the ITG, more posterior to non-math sites in the SPL, SFG, and mSFG (although the correlations for SFG and mSFG did not survive the FDR correction for multiple comparisons); more anterior to non-math sites in the IPS, MFG and IFG, and more dorsal to non-math sites in the SMG (**Figure 2**).

### Profile of Activity Across Anatomical Regions

By leveraging the temporal resolution and the high signal-to-noise ratio of the iEEG signal, we explored the pattern of electrophysiological changes within a given neuronal population from the time of stimulus onset to the time of subject’s responses. During Task I, when arithmetic equations were presented at once, sites with preferential responses during arithmetic condition exhibited a similar profile of sustained responses. During Task II, in which the arithmetic stimuli were presented sequentially, we discovered a stereotyped signature of activation across sites. Neuronal population responses in math preferential sites were significantly higher after operand and response conditions (e.g., “2” and “3” and “5” in “2 + 3 = 5”) as compared to arithmetic operation symbols (e.g., “+” and “=” in “2 + 3 = 5”). Interestingly, the activity induced by the second operand was also significantly higher than the activity induced by the first operand. This is noteworthy because after the first operand and first operation symbol (e.g., “2” and “+” = in the above example), the brain is yet to be engaged in arithmetic processing. The brain starts calculating when the second operand (in this case “3”) is shown. The profile of activity in math preferential sites was in sharp contrast to the profile of non-preferential sites such as early visual areas or sensory motor cortices (**Figure 3**).

In parallel to activations during arithmetic processing, we found sites that showed clear decreased activity during the math condition (n = 275 across 46 subjects). That is, HFB activity was reduced significantly below the baseline period, in both Simultaneous and Sequential Calculation tasks (**Figure 4**). The deactivated sites were also anatomically consistent across subjects and located in the posteromedial cortex (PMC, n=36, 13.4%), medial temporal gyrus (MTG, n=37, 13.4%), orbitofrontal cortex (OFC, n=20, 7.2%), SFG (n=15, 5.4%), superior temporal sulcus (STS, n=13, 4.7%) and the medial prefrontal cortex (mPFC, n=13, 4.7%) – all of which are known to be regions of the default mode network (57). Importantly, regions with decreased activity for math also showed the strongest signal decreases after the second operand at the same time that the math-preferential sites showed the highest activations – possibly suggesting a synchronous antagonistic relationship across the math preferential and math deactivated sites.

### Temporal Order

We calculated the concentration of HFB signal throughout each trial for each site with preferential math responses (n=248, across 48 subjects) in the math condition of the Simultaneous Calculation task. Since the task was self-paced and trials had varying RTs within and across subjects, we normalized the HFB power by the % RT at each single trial (i.e., calculating the HFB power in the 0-100% time-window of each trial). By comparing the temporal profiles of average HFB concentration across sites (**Figure 5B**), a clear order of activation emerged: sites were “ignited” in a successive order. Interestingly, the earlier activations did not cease when the next batch of sites were “turned on”. The earlier sites continued their activity simultaneously with the later ones until all sites ended their activity as the subject responded (**Figure 5B and C**).

**Figure 5.**
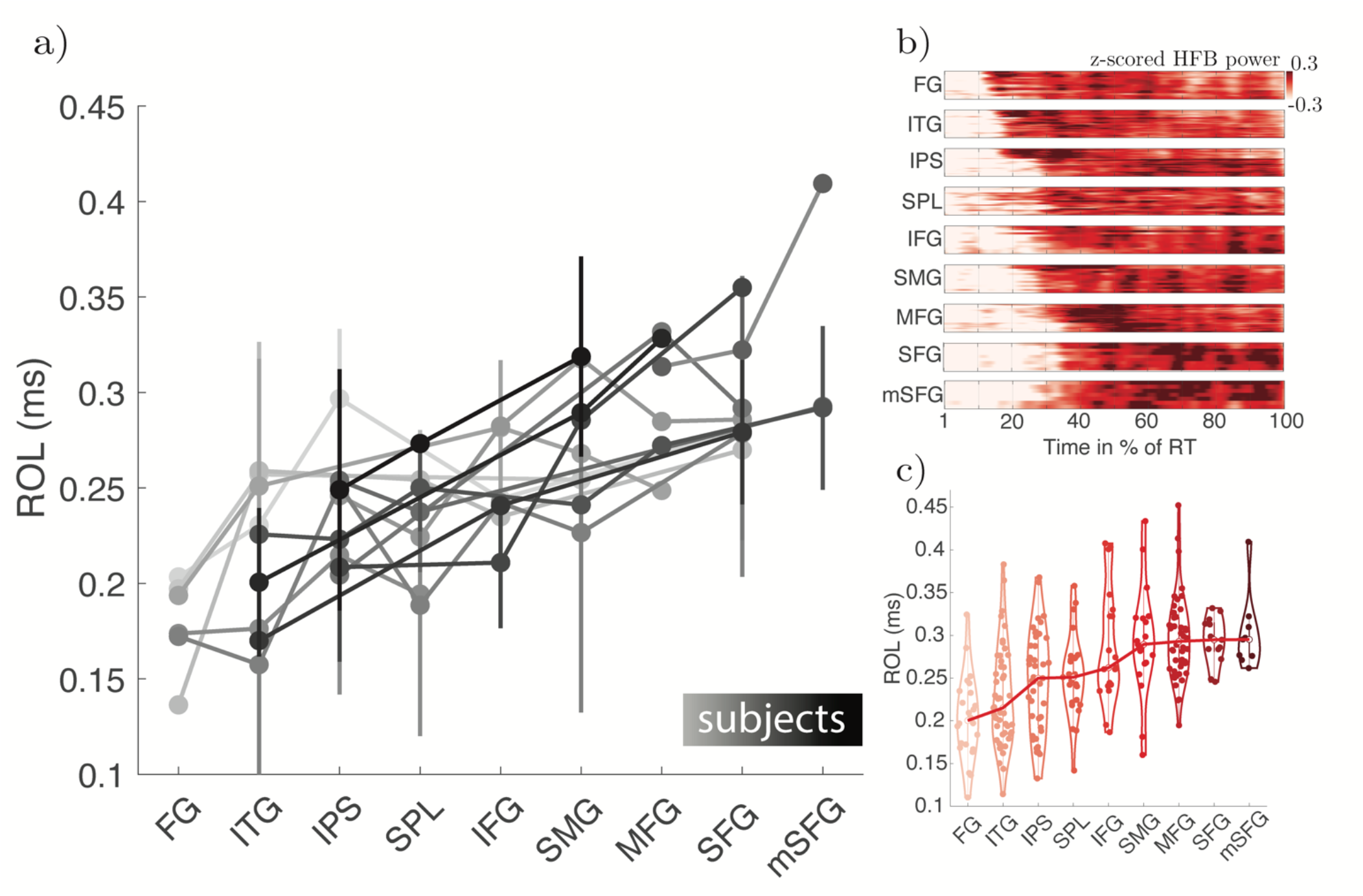
Temporal dynamics of math processing across regions within individual brains. A) In 16 subjects, we had simultaneous coverage in at least 3 of the 9 ROIs. Each line (with a shade of gray) represents a single subject. While subjects had dissimilar coverage across the 9 ROIs, it is noteworthy that the slope of the linear regression is preserved for all subjects - independently of the particular group of ROIs covered in each individual brain. Error bars represent the standard error of the mean. B) RTs were normalized across subjects, and the power of high-frequency broadband (HFB) was plotted for each region. Each row within each matrix represents the averaged signal across trials of individual electrode sites within each anatomical region. These sites have been sorted according to their average ROL. One can clearly see a correspondent progression of time of onset of HFB activity (i.e., response onset latencies, ROLs) across regions. C) Distribution of ROL values recorded for each of the 9 ROIs across subjects is shown in violin plots with dots representing an averaged ROL of a single electrode across all math trials. See Video 1.

Next, we measured the precise time of onset of HFB activations (Response Onset Latency, ROL) at each site. At the group level, we observed a very clear linear trend. Activations start in the temporal lobe sites and then parietal sites before the frontal areas are engaged (**Figure 5B and C**). This was confirmed with an ANOVA for linear regression model (F(8,239) = 11.192, R2 = 0.27, p < 0.001). In this analysis, we sought to statistically demonstrate a linear trend in the data after sorting the regions by mean ROL. Note that the purpose of this analysis was not to establish the order itself (which would be circular), but rather to confirm the presence of a statistically significant linear trend in the sorted data. To verify this finding at the single subject level, we found 16 subjects who had simultaneous recordings in at least 3 of the ROIs. For each subject, we first re-sorted the ROIs by excluding their individual data from the group mean. This procedure ensured that there was no circularity in this analysis. Out of the 15 subjects, only one exhibited a change in the order, specifically with mSFG preceding SFG. As a result, we chose to exclude this subject from subsequent analyses and from Figure 5A. Next, we fit a linear regression model on the ROLs as a function of sorted ROIs. We found that the slopes of temporal order within each individual were all positive and remarkably consistent (mean = 13.3 ms, std = 11.3 ms, R^2^ of the linear fit ranged from 0.35-0.99, mean=0.66, sd=0.24) irrespective of the particular set of simultaneous ROIs recorded (**Figure 5A**).

In order to probe if the temporal sequence of activations could be captured at the single trial level, which would reflect coordinated activity across regions, we selected a single subject who had simultaneous coverage in not only ITG, IPS and mSFG, but also in early visual areas (lateral occipital gyrus (LOG), posterior SPL as well as the precentral gyrus (PCG). For each electrode pair, we ran a cross correlation analysis and estimated the indices of functional connectivity (*Δr_peak_*) and temporal delay (*Δr_lag_)* (see Methods). If two regions are synchronized, the *Δr_peak_* should be high and positive, and if one region A is leading and another region B is lagging, *Δr_lag_* should be positive. As seen in **Figure 6A**, the early visual region (LOG), despite being the first in the cascade of activations, and despite showing a significantly high initial response for arithmetic stimuli, does not show synchronized responses with downstream regions. However, the arithmetic ROI sites of ITG, IPS and mSFG show strong evidence of synchronization with each other. Activity in these sites also correlated with the activity of subsequent regions in the processing chain even if their activity was later and aligned with the time of the subject’s response (i.e., anterior SPL and the precentral gyrus in the motor cortex). These correlations clearly show a progressive lag following the processing chain order as revealed by the ROL analysis. Interestingly, we observed decreased functional connectivity across pairs of brain regions as their temporal lag widened. In other words, as the temporal lag and order between two regions increases, their functional connectivity decreased: ITG with - IPS (*Δr_lag_* = 4 ms; *Δr_peak_ = .*20), mSFG (*Δr_lag_* = 70 ms; *Δr_peak_ = .*15) pSPL (*Δr_lag_* = 490 ms; *Δr_peak_ =* 0.07) and PCG (*Δr_lag_* = 788 ms; *Δr_peak_ =* 0.06). IPS with mSFG (*Δr_lag_* = 36 ms; *Δr_peak_ =* -.18) pSPL (*Δr_lag_* = 486 ms; *Δr_peak_ =* 0.11) and PCG (*Δr_lag_* = 1.01 ms; *Δr_peak_ =* 0.07). mSFG with pSPL (*Δr_lag_* = 332 ms; *Δr_peak_ =* 0.11) and PCG (*Δr_lag_* = 920 ms; *Δr_peak_ =* 0.09); and pSPL-PCG (*Δr_lag_* = 436 ms; *Δr_peak_ =* 0.18). To replicate what we had observed in the exemplar single subject, and to probe if the pattern of electrophysiological correlation was reproducible across subjects, we calculated the degree of cross-correlation across pairs of electrodes located between any of the 9 regions of interest. We found 219 pairs of electrodes with math-preferential responses in 29 subjects who had simultaneous recordings across some (but not all) nine regions of interest. Note that we used all pairs of electrodes to maximize the dataset, so for example if a given subject had two electrodes in the IPS and one in ITG, we included the pairs IPS_1-ITG and IPS_2-ITG. Next, we plotted the degree of correlation for a given ROI to all the other ROIs ordered by temporal distance based on the response onset latency measures. This analysis revealed that for most ROIs (6 out of 9) there was a significantly stronger correlation between pairs of electrodes that were temporally adjacent, and the strength of connectivity decreased with temporal distance across pairs of regions. In two (IFG and SMG) regions in which this tendency did not reach statistical significance, the correlation coefficient was also negative, suggesting a similar, but weaker effect.

**Figure 6.**
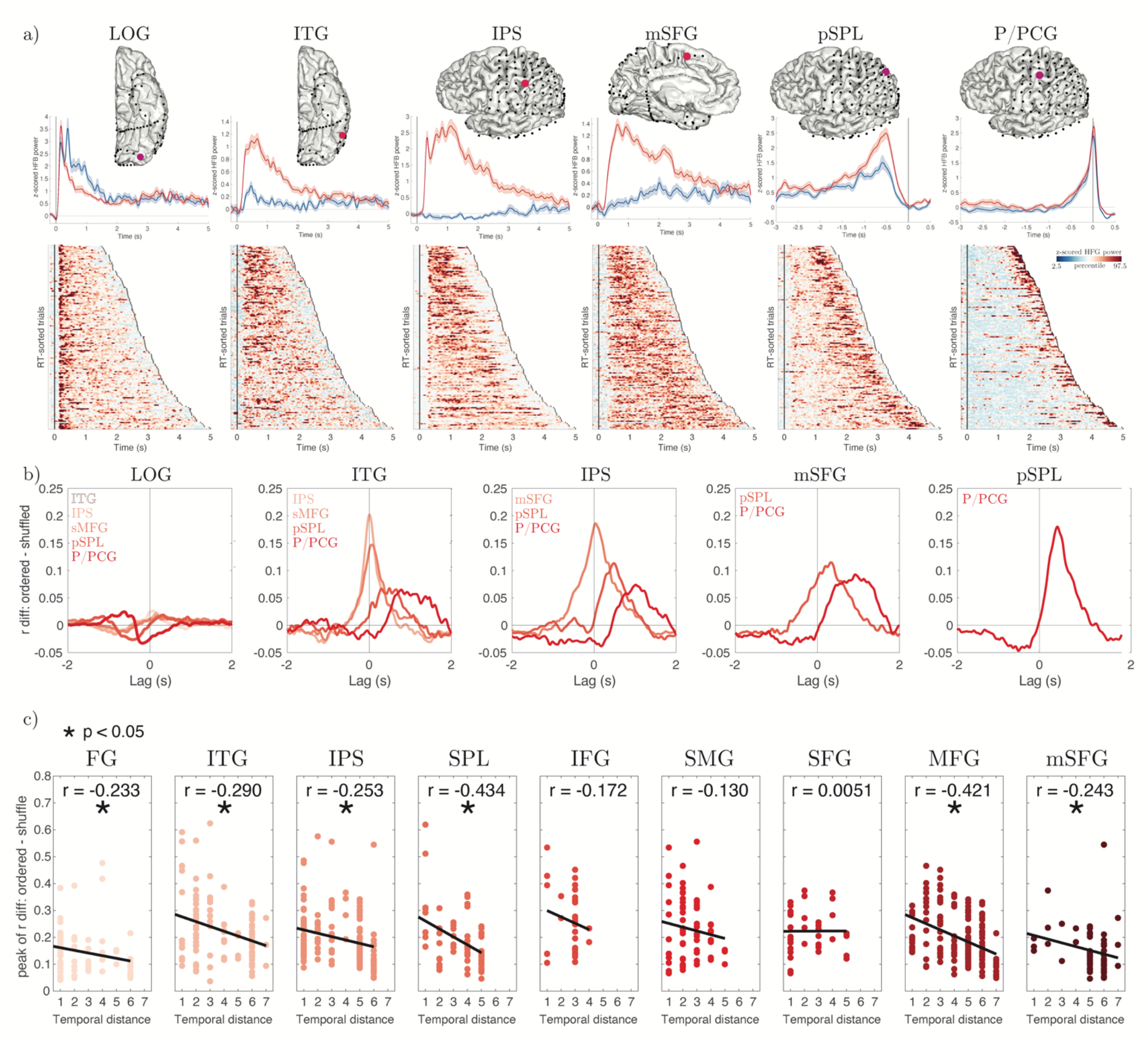
Functional connectivity of the main math ROIs. A) Exemplar subject with simultaneous coverage of 5 regions activated during math, from earlier visual to the motor cortex. ITG, IPS and mSFG showed strong math selectivity. Top time courses show averaged activity across trials for math and memory. Bottom plots show single trial activity ordered by RT (trial ends when subject presses the response button). Shaded error bars represent the standard error of the mean across electrodes. B) Same exemplar subject: lagged correlation between the activated regions. Lines represent the pairwise correlation coefficient with preserved trial order subtracted from the average correlation from the distribution of 2,000 pairwise correlations generated by shuffling the trial order. The difference between correlation values from ordered and shuffled data inform how the regions are functionally connected at the single-trial level. Rightward peaks mean that correlations are delayed. C) Electrophysiological functional connectivity based on lagged-correlation for each math preferential/selective ROI with all the other regions, grouped using the temporal distance between them, calculated from the ROL analysis.

### Exploring Feature Dependent Change of Activity

Although our experiments were *not* designed to decipher the contribution of each brain region in specific cognitive aspects of arithmetic processing (i.e., which region is contributing to what), we performed an exploratory analysis as an attempt to model the activity of each math-active site with a stepwise multiple linear regression - using the HFB activity as the dependent variable and several arithmetic “features” as predictors. We only used data from the Sequential Calculation task because, by design, this task isolates two distinct moments of interest: the calculation stage (following the presentation of the second operand) and the verification stage (following the presentation of the proposed result). For instance in 3 + 4 = 8, the calculation stage only begins after the second operand “4” is shown and the verification process only begins after the result “8” is presented.

We explored the effects of 5 main arithmetic features that had significant impact on the measure of RT. These can be seen as indirect indices of “*problem difficulty*”: min operand, max operand, cross-decade, format and absolute deviant (see *Behavioral Performance* section). In brief, *min operand* is the smallest operand, the *max operand* is the largest operand, *cross-decade* means that the product of a calculation is above the decade of max operand, *format* refers to digit vs. number word, and *absolute deviant* is the difference between the correct answer and the one presented in the equation.

During the calculation stage, we found that the HFB power in 55 recording sites was modulated by the min operand effect and the majority of these sites (64%) also displayed preferential responses to math compared to non-math condition (**Figure 7**). Max operand and cross decade features also modulated the HFB responses in several sites, with the majority of sites showing higher HFB activity with higher *min operand* (78%), *max operand* (92%), and equations with a *decade cross* (72%). Only a small proportion of these sites showed preferential responses for math. The format modulated 93 sites but only a minority (31%) showed preferential responses for math, suggesting that majority of math preferential sites are format independent and that other areas related to visual processing and language contain a large number of sites modulated by digits vs. number words, which might have little to do with calculation *per se*. Among these sites, 61% had higher activation for number words compared to digits. Finally, as expected, no site showed significant modulation by the absolute deviant, since that information is not available at the calculation stage. During the verification stage, 55 sites (44% math preferential) were modulated by the absolute deviant, with the majority responding stronger for increased magnitude of the deviant (64%).

**Figure 7.**
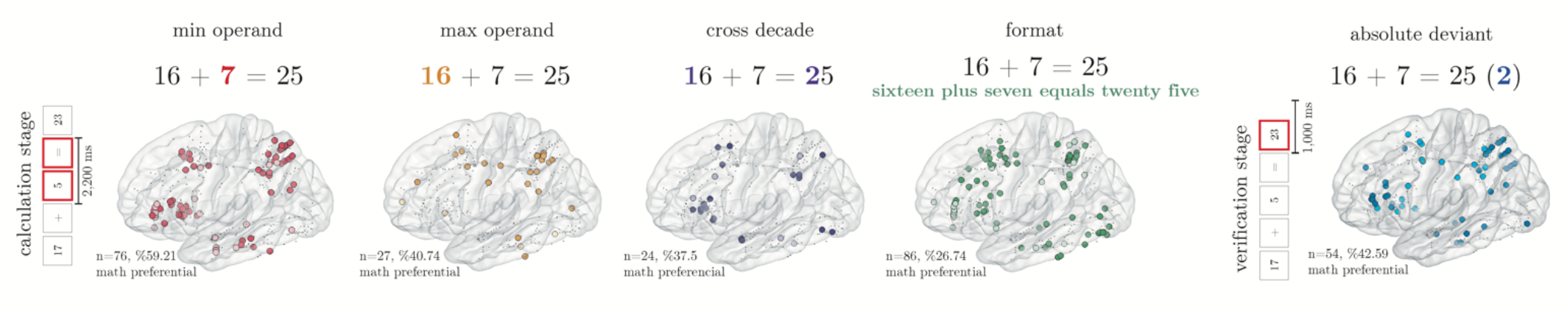
Encoding of arithmetic features across the math active sites during the Sequential Arithmetic task. Large and colored dots represent sites in which a given feature significantly explained variance of the HFB activity (after second operand) and therefore were retained by the stepwise multiple regression fit. The darker color for min operand, max operand and absolute deviant represents sites in which the HFB activity increased with the magnitude of these features, whereas the lighter color represents sites in which the HFB activity decreased with the magnitude of these features. For cross decade, darker color represents sites in which the HFB activity was stronger for problems that crossed a decade (sites with the opposite effect are in showed in lighter color). For format, darker color represents sites in which the HFB activity was stronger for number words as compared to digits (sites with the opposite effect are in showed in lighter color).

## DISCUSSION

Our present study complements the prior imaging studies (19, 24, 29, 52, 58, 59) by providing hitherto unknown information about the *timing* of events in the brain during arithmetic processing. We recorded from a large number of sites in a large number of human subjects while they performed two arithmetic tasks. We validated data between the two tasks and across individual subjects with recordings in the same regions of the brain. We then characterized, for the first time, the precise temporal dynamics of distributed but anatomically precise responses across co-activated neuronal populations in the human brain. This novel information will advance our understanding of the timing of events across multiple regions of the brain and will addresses some of the key open questions as detailed below.

### Activity of a very large mantle of the human brain is changed when we engage in simple arithmetic

Subtractive approaches often used in neuroimaging methods contrast responses during a target condition with responses during the non-math condition. This subtractive method often segregates different regions of the brain and makes it impossible to differentiate conditions that lead to similar changes in activity from conditions that lead to no change in activity. Here, we compared the activity of specific populations of neurons (i.e., sites of recordings) during the math condition compared to their *own activity a few hundred milliseconds prior to the onset of the math condition* (i.e., baseline). We observed 45% of all recording sites across 85 subjects showing significant activity. The engagement was distributed across virtually all sampled brain regions and in both hemispheres and, interestingly, while 80% of the involved sites showed increased activity during arithmetic tasks compared to the baseline period, 20% displayed decreased activity. Our results demonstrate that arithmetic processing is highly distributed in the human brain, which better aligns with a more recent understanding of brain functions (60) and corroborates many prior observations with non-invasive neuroimaging studies suggesting a widespread activation of the brain if one considers the baseline activity (rather than a different cognitive function) as a control condition. These findings impose constraints on localizationist models of mathematical processing that remain the most influential in the literature (20, 29, 61). Our results suggest a broader landscape of brain regions participating in mathematical cognition than previously acknowledged. This includes not only areas demonstrating activation in response to mathematical stimuli, but also regions manifesting deactivation. This diverse array of responses underscores the complexity and intricacy of the brain’s processing mechanisms for mathematics, suggesting that our understanding of these processes needs to consider a more holistic and interconnected network of regions rather than isolated localized areas.

### Math-preferential and non-math-preferential sites are orderly co-localized

To make a bridge to the neuroimaging literature, we then employed a subtractive method (akin to the ones used in neuroimaging studies) and found nine regions with significantly higher responses during the target (i.e., math) condition. The anatomical location of these sites was remarkably consistent across subjects and replicated the findings reported in previous fMRI studies (29), namely: fusiform gyrus (FG), inferior temporal gyrus (ITG), intraparietal sulcus (IPS), superior parietal lobule (SPL), supramarginal gyrus, inferior (IFG), middle (MFG) and superior frontal gyrus (SFG), and medial superior frontal (mSFG) gyri. In our analysis, we defined the preferentiality of responses by the presence of significant activation during math condition and *no activations* during the control condition. However, we emphasize that the set of regions showing preferential responses during the math condition may not represent a “*selective”* math network as their preferential response during the math condition compared to non-math condition does not imply that they are *solely* involved in arithmetic processing. In keeping with this, we recently showed that electrodes in the posterior parietal cortex with preferential responses to math also showed preferential responses to visuospatial cues appearing in the contralateral, but not ipsilateral, visual fields (23). Furthermore, as we discuss later, our multiple regression analysis revealed that the encoding of different arithmetic features is distributed across brain regions and is not limited to the sites with strongest math-preferential responses.

A potential confound for the selectivity analysis is the related to possibility that the math condition may have generally been more demanding for some subjects. Nevertheless, we posit that the task demand isn’t the solitary determinant and present two reasons to substantiate this claim. First, as elucidated in our reaction time analyses, an interesting pattern emerged. We observed higher RT for the math condition when compared to the non-math condition in the simultaneous task. Yet, a reverse pattern was notable in the Sequential task, where the non-math condition yielded higher RT in comparison to the math condition. Secondly, our approach to identifying a site as “math preferential” was predicated on three specific criteria. The site needed to exhibit significantly higher HFB activity during math trials as compared to baseline, and higher trial HFB activity during the math condition in relation to the non-math condition. Of paramount importance, though, was the third criterion: the site was not to show a significant difference in trial HFB activity during the non-math condition when compared to baseline. The third criterion is particularly salient, given that task demand was also a component in the non-math (memory) condition, albeit to a lesser extent. Rather than reporting a “graded” response, we observed that the math preferential sites demonstrated no responses during the non-math condition. This rigorous definition bolsters our belief that the observed preferential responses are, in large measure, driven by the task content.

Neuroimaging studies often highlight a set of brain regions with increased hemodynamic responses during a target condition. Such group-based findings may be misinterpreted as if the entire region is homogenously activated during the target-condition across individual brains. Our findings provided a glimpse of functional organization in the human brain in the mesoscale (millimeter) level. Neuronal populations with increased activations during the target math condition were found to be co-localized adjacent to populations with preferential responses during the non-target (memory) condition, and that neuronal populations with opposite profiles of activity were orderly co-localized, i.e., anterior proclivity for math-preferentiality was seen in IPS, MFG, and IFG; posterior proclivity in SPL and SFG; dorsal proclivity in the SMG; and right hemispheric proclivity in the FG. In other words, adjacent populations of neurons that are anatomically juxtaposed with each other may have preferentiality for different cognitive functions. That might explain why neuropsychological patients almost always display a variety of cognitive deficits spanning different cognitive functions, following a lesion or atrophy in specific brain regions (34, 35). Our results indicate that some brain regions may indeed be engaged in both mathematical and linguistic processes and that these cognitive symbolic systems could share some underlying computational mechanisms. This perspective aligns with a growing body of evidence pointing towards the overlap in neurodevelopmental disorders, reinforcing the idea that these disorders are complex, multifactorial entities that often manifest in a spectrum of probabilistic subtypes (62). It is noteworthy that estimates suggest up to 40% of children with dyscalculia (mathematical deficits) also exhibit symptoms of dyslexia (reading deficits) (63). Therefore, our findings underscore the need to move beyond a compartmentalized view of cognitive functions and towards a more integrated understanding of brain activity.

Although our study did not specifically formulate hypotheses regarding the within-region anatomical organization of juxtaposed activations, we believe our findings provide a robust foundation for future research. This body of work could delve deeper into the intricate anatomical structuring associated with high-level cognitive functions, thereby advancing our understanding of cerebral organization.

### Successive engagement and recurrent interactions

One of the most important findings in our study pertains to our observation of temporally ordered successive “ignition” of brain regions during simple arithmetic function at the single-subject level. As noted, a much larger extent of the brain was activated during the math condition. However, by focusing on the select sites of interest (i.e., nine anatomical regions with math-preferential responses), we explored the temporal order of activation across regions and found a successive order of activations, *within the sub-second space,* from FG, to ITG, IPS, SPL, IFG, SMG, MFG, lateral SFG and lastly to the medial-SFG sites (**Figure 5** **B and C**). More importantly, distinct regions of the brain became engaged and remained engaged above their own baseline across the length of each trial until the subject responded. It should be noted that our finding of temporal order of activations was not an artifact of averaging across different brains. As clearly documented in **Figure 4**, in 16 subjects with simultaneous recording across several of the same ROIs, a similar temporal order and slope was observed. Across subjects, the coefficient of this temporal order was nearly identical even though each subject performed the task with variable RTs.

Our observations from the cross-correlational analysis showed that the temporal precedence of activation was not sufficient to influence the activity profile of the rest of the chain of regions. Instead, regions that are temporally closer to each other seem to be more strongly influencing each other at the single-trial level. As regions get further apart in temporal order, their degree of influence decreases. For instance, in an exemplar subject with simultaneous coverage in multiple math-preferential regions, we found a very weak degree of connectivity between the first and the rest of the regions that responded. However, we observed a strong functional relationship (with an expected temporal lag) between the ITG and IPS.

Although the aim of the current study was not to arbitrate between different architectures of information processing in the brain, our real-brain-data can be taken as a support of previously proposed theoretical models such as the ordered and functionally dependent cascade of partially overlapping brain states (64). Our observation that the information gets progressively lost or transformed along the processing chain also supports theoretical models such as leaky competing accumulator model (65). The notion of “information leak” in cognitive processing alludes to the imperfect retention or utilization of information essential for task performance at each step in the processing chain. This may occur due to either the imperfect transmission of information from one brain region to another, or because a brain region does not fully utilize the information it receives. Several factors could contribute to this “leakage”, such as inherent noise in neural signaling, the complexity of the information being processed, or limitations in the brain’s computational capacity (66). From a neurophysiological perspective, these information leaks or transformations could manifest through various mechanisms. For instance, the decay or dispersion of the neural signal as it travels between brain regions, or the selective filtering of information based on task relevance (67). Moreover, the transformation of information could refer to how information is repackaged or reformatted across different regions or processing stages. Initial processing stages may involve encoding basic numerical data or mathematical operations. In contrast, later stages could involve more complex processes like error checking, problem-solving strategy selection, or the integration of numerical data with spatial, temporal, or other contextual information. Consequently, the same piece of information could take on different forms or representations as it progresses along the processing chain.

Overall, our findings are in line with a growing literature that proposes that even simple tasks like object recognition (68), visibility judgment (69) and hierarchical perceptual decisions (70) involve a distributed network of recurrent interactions, beyond purely feedforward architectures (71). Our work expands on these findings from very simple operations and suggests that a similar architecture underlies the processing of complex human unique abilities such as symbolic arithmetic.

### Relating the spatiotemporal dynamics of brain activity to specific stages of cognitive processing

While the current study was not designed to address the neural mechanisms underlying specific stages of cognitive processing during the target condition (i.e., mathematical processing), our data revealed important layers of information that can guide future mechanistic explorations. For instance, we observed a clear profile of activations with higher HFB activity primarily following the second operand and after the presentation of the equation results. These time points are precisely the times in which subjects become engaged in arithmetic calculation and comparison between the correct vs. presented results. Interestingly, in the sites deactivated during the arithmetic condition, we discovered that the onset of such decreased activity occurred after the presentation of the second operand matching (and coinciding in time with) the sharp activations in the functionally opposing regions. These observations were remarkably reproducible across subjects and consistent with our own preliminary findings in a few regions of the brain(31, 37, 72). Moreover, by manipulating the format of presentation (digits. vs. number words), we showed that the majority of the math preferential sites are agnostic about the specific format in which the numbers are presented, and therefore operate on an abstract level.

We should clarify that our current tasks were not specifically tailored to examine the discrete role of each brain region during mental calculations. Therefore, our findings should be viewed as an initial foray towards a comprehensive and precise spatiotemporal dissection of the computations performed at individual brain sites during arithmetic processing. One noteworthy finding from our multiple regression analysis suggests a distributed encoding of different cognitive features, challenging the notion that these features are confined solely to regions displaying preferential responses during the target condition.

These observations don’t just question prevailing notions of brain functional organization; they also challenge established models of arithmetic processing, such as Dehaene and colleagues’ Triple Code Model (61) and others (29). In these models, specific functions related to numerical and arithmetic processing are assigned to each area within the potential math network. However, our study unveils a more distributed pattern of brain region engagement during mental calculations, which does not align perfectly with the functional compartmentalization proposed by models like the Triple Code Model.

Our results thus call for a re-evaluation of traditional models of arithmetic processing and underscore the necessity for further research incorporating a more distributed conception of brain function. Future iEEG studies, with targeted experimental tasks, ought to employ a whole brain level of analysis to decipher the contribution of each brain region of interest in specific features of the arithmetic condition.

### Closing remarks

Our large-scale anatomical, temporal and functional iEEG findings provide novel information about the profile of neural activity across different regions of the human brain to help characterize how different regions of the brain work collectively in the service of a given task. As such, the present work ought to be taken as a prototype model for how different anatomical populations in distinct regions of the brain change their activity in unison with each other to enable a human subject to perform a cognitive task as simple as 2 + 2 = 4. We hope that our findings, along with the acquired raw data, which will be shared in a public repository, can be used as real brain data to construct more accurate theoretical models of distributed brain activity.

## Figure and video legends

**Video 1**. Spatiotemporal activations of the Simultaneous Calculation task. In the Simultaneous Calculation task, subjects judged the accuracy of full arithmetic equations presented in X + Y = Z format (e.g., 16 + 8 = 22). This video show how the power of activity in the high frequency broadband (HFB) changes from ∼100ms before until ∼1000ms after the stimulus onset in select electrodes with preferential activity during the math condition.

**Video 2.** Spatiotemporal activations of the Sequential Calculation task. Same as Video 1, but the responses are measured during the Sequential Calculation task, when subjects viewed math stimuli appearing sequentially one at a time in “X”, “+”, “Y”, “=”, “Z” format (e.g., “16”, “+”, “8”, “=”, “22”).

## Extended data legends - The full Extended data can be found after the references

Figure 1-1. Subject’s demographics, basic neuropsychological information and task completion.

Figure 1-2. Stimuli list for the Simultaneous Calculation task

Figure 1-3. Stimuli list for the Sequential Calculation task

Figure 1-4. Math preferential sites per anatomical region

Figure 1-5. Non-math preferential sites per anatomical region

Figure 1-6. Math and Non-math equally preferential sites per anatomical region

Figure 1-7. Math deactivated sites per anatomical region combining the Simultaneous and Sequential calculation tasks results.

Figure 1-8. Arithmetic problem-size effect by subject. Supports Figure 1.

## Data sharing plans

The dataset will be shared upon request. All code to preprocess, analyze and visualize the data is available at: https://github.com/LBCN-Stanford/lbcn_preproc

**The authors declare no conflict of interest associated with this manuscript.**

## EXTENDED DATA COMBINED

**Figure 1-1.**
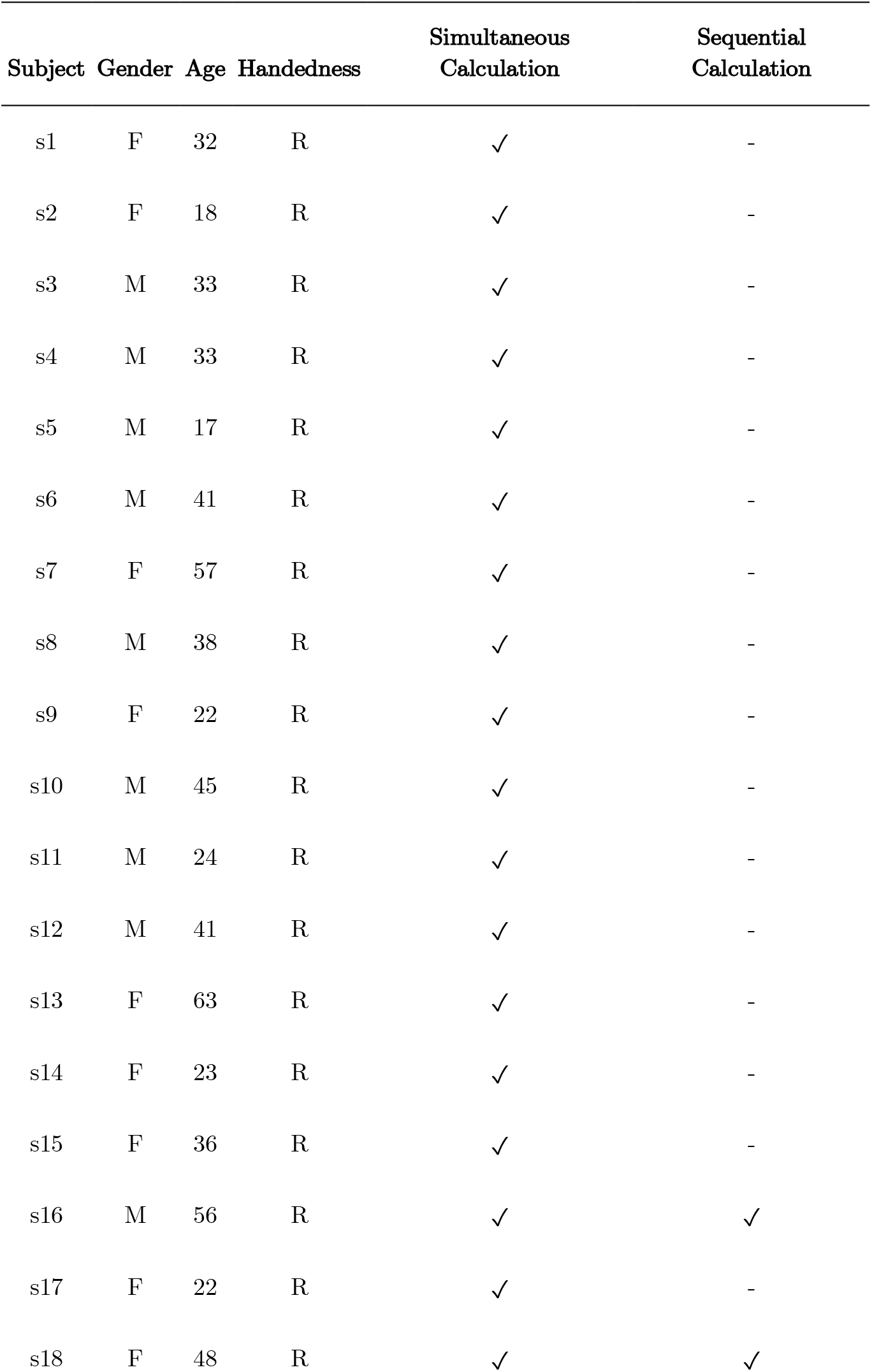
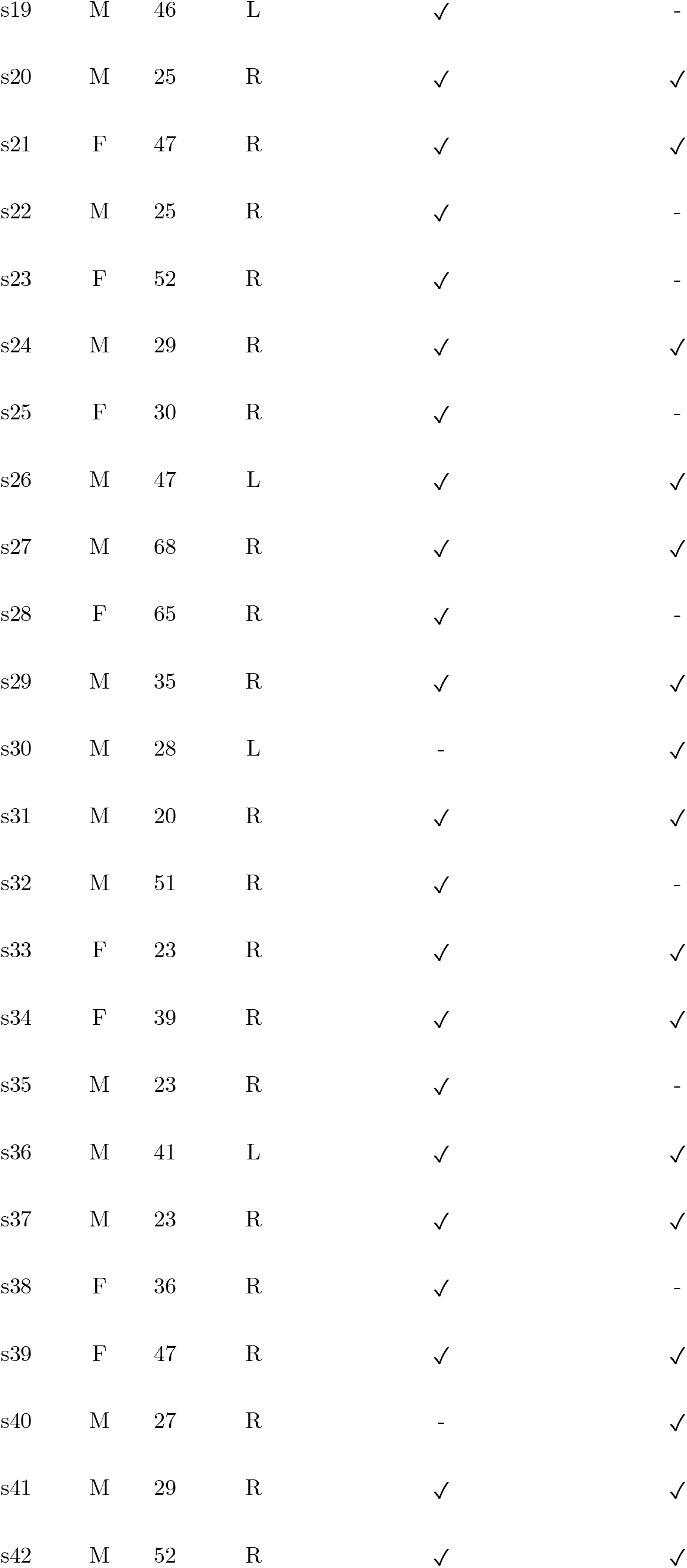
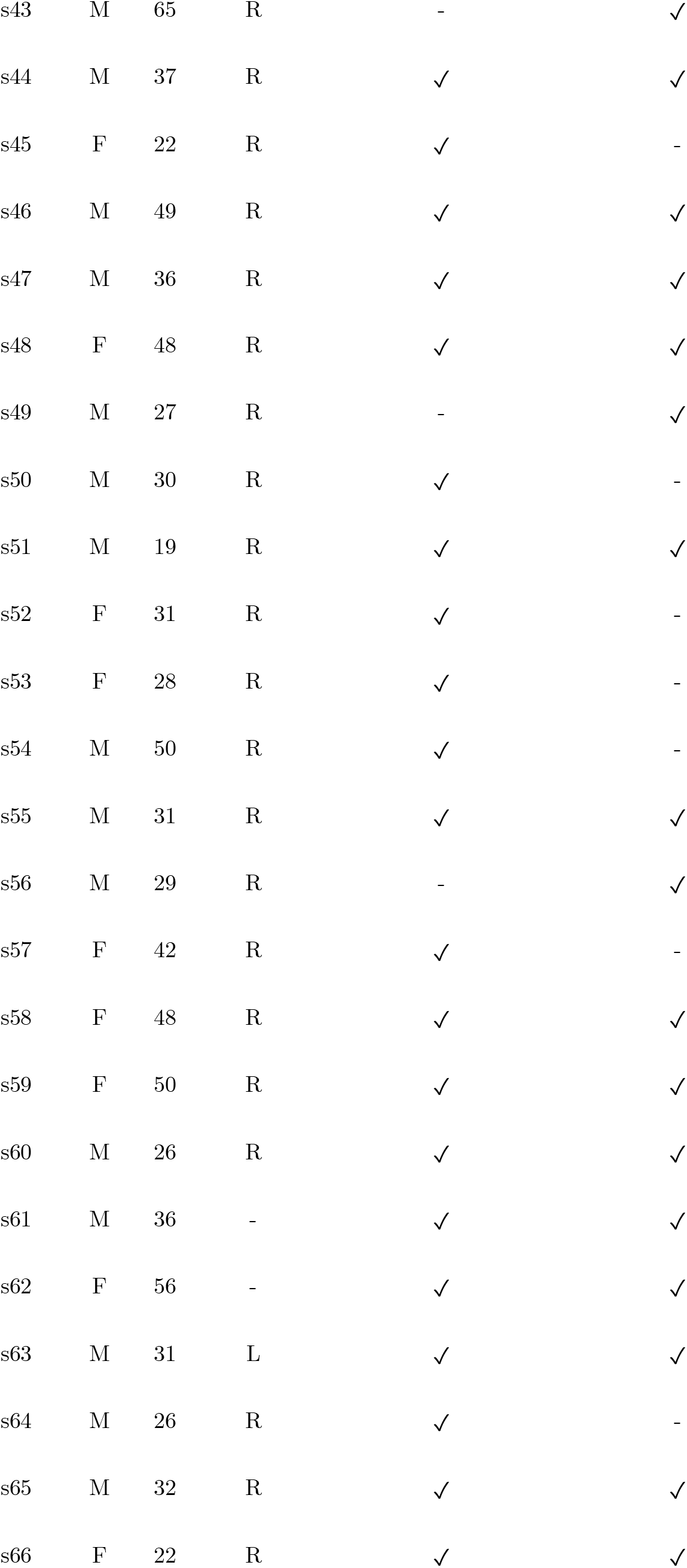
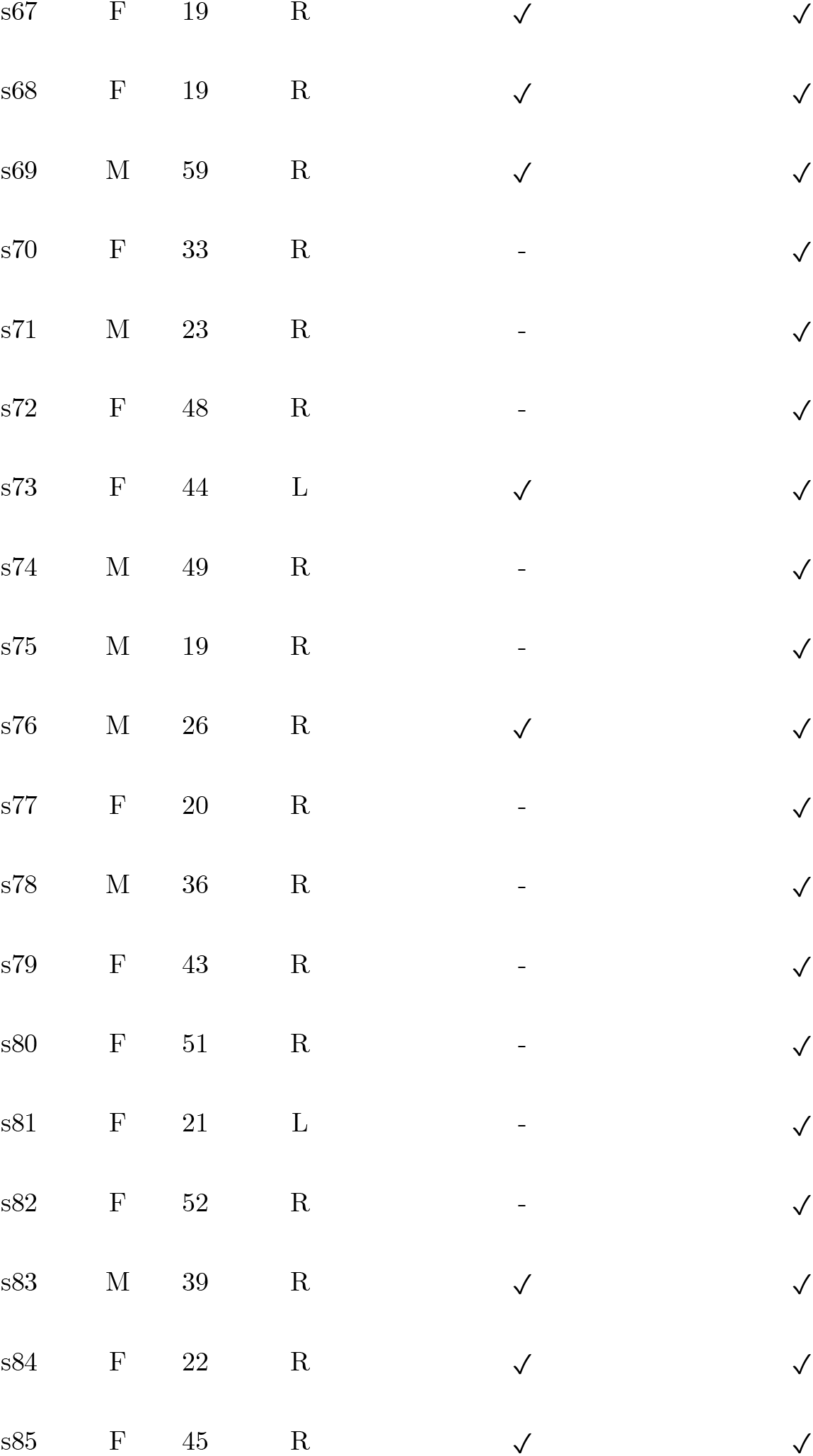
Subject’s demographics, basic neuropsychological information and task completion.

**Figure 1-2.**
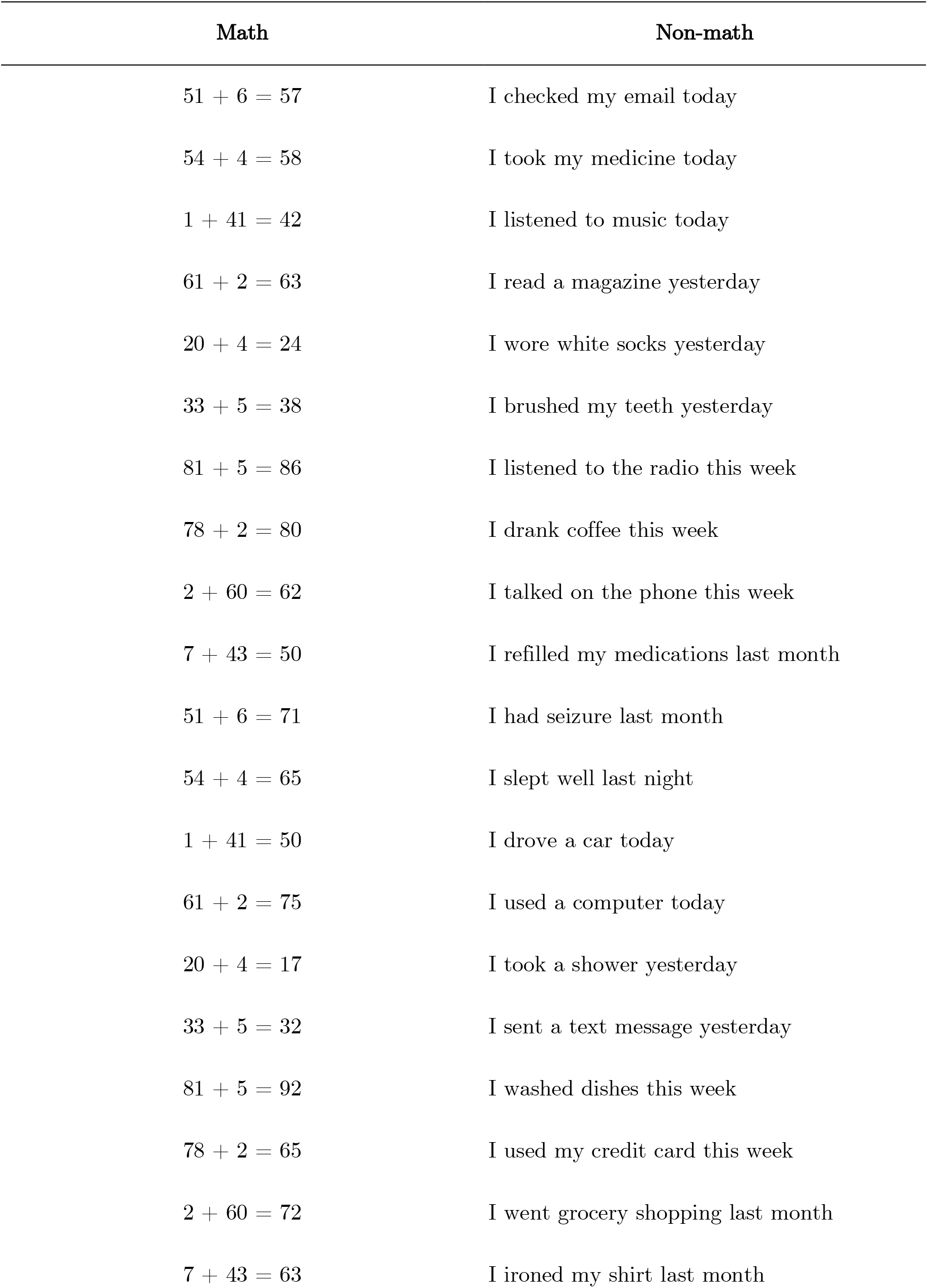
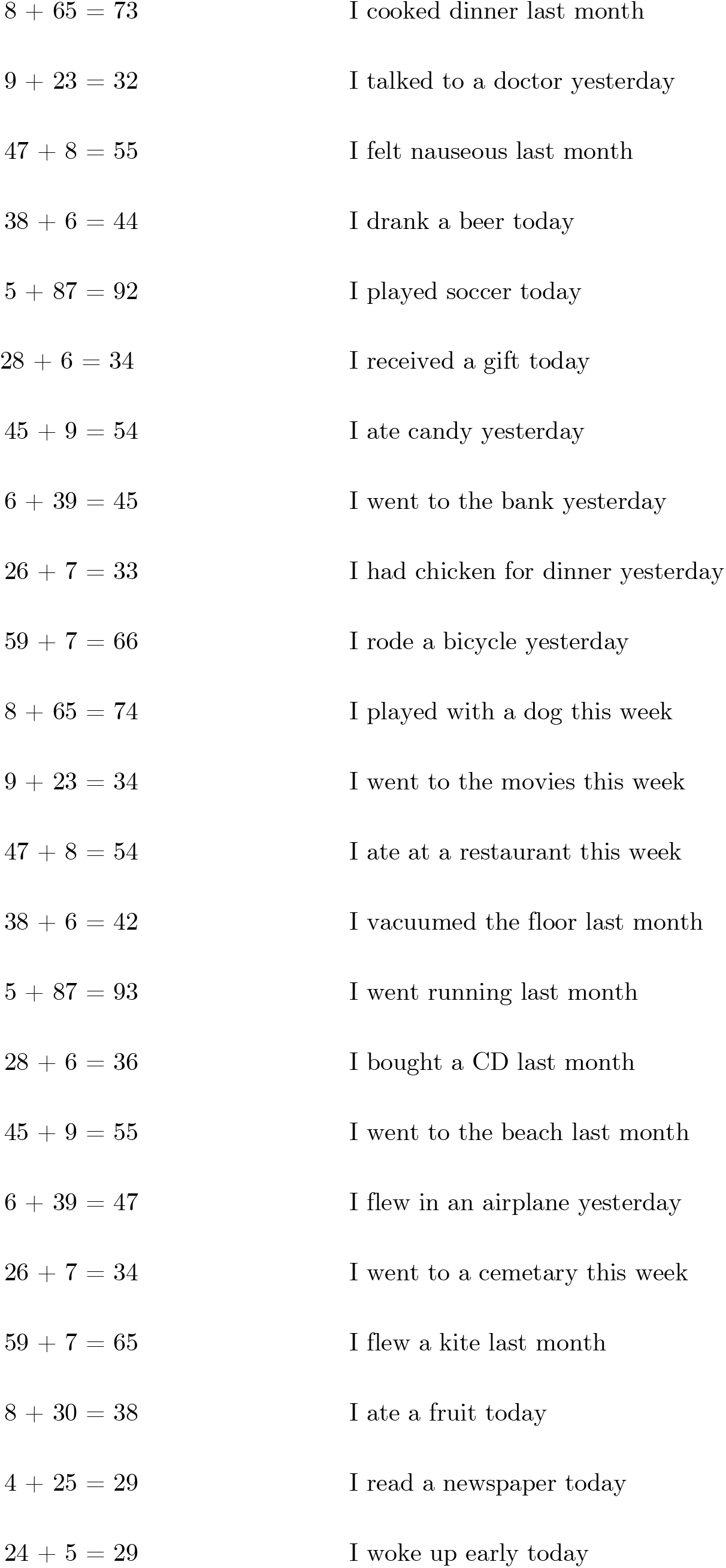
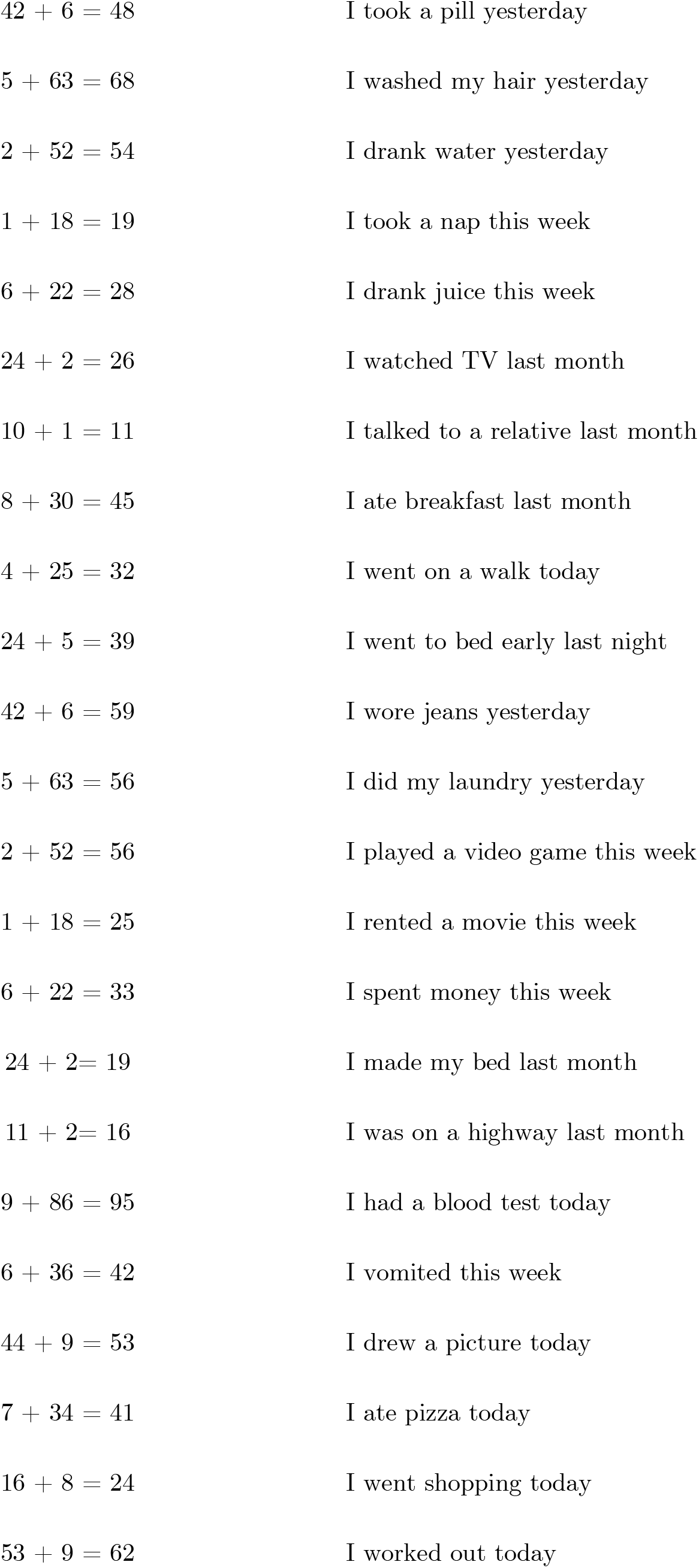
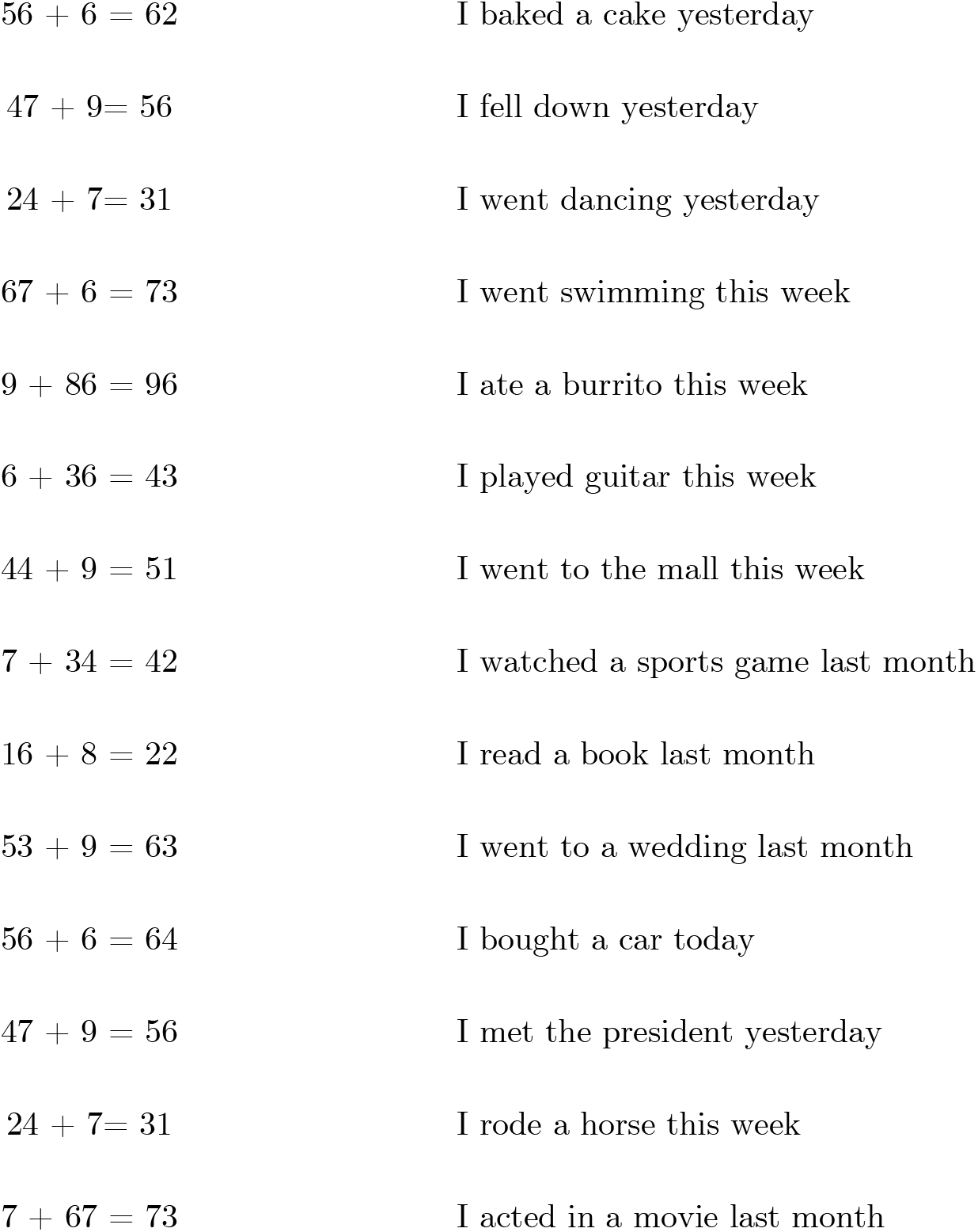
Stimuli list for the Simultaneous Calculation task.

**Figure 1-3.**
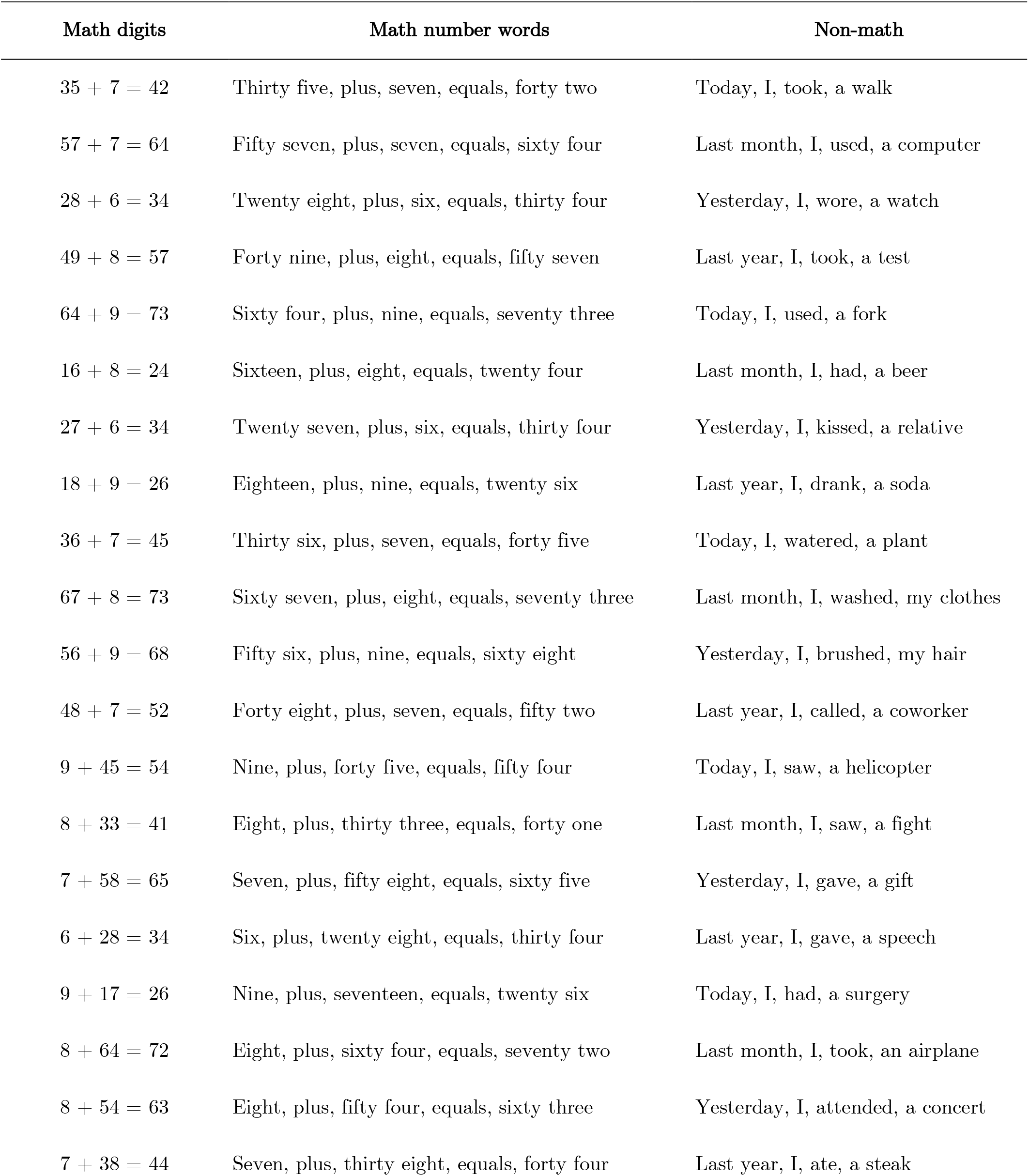
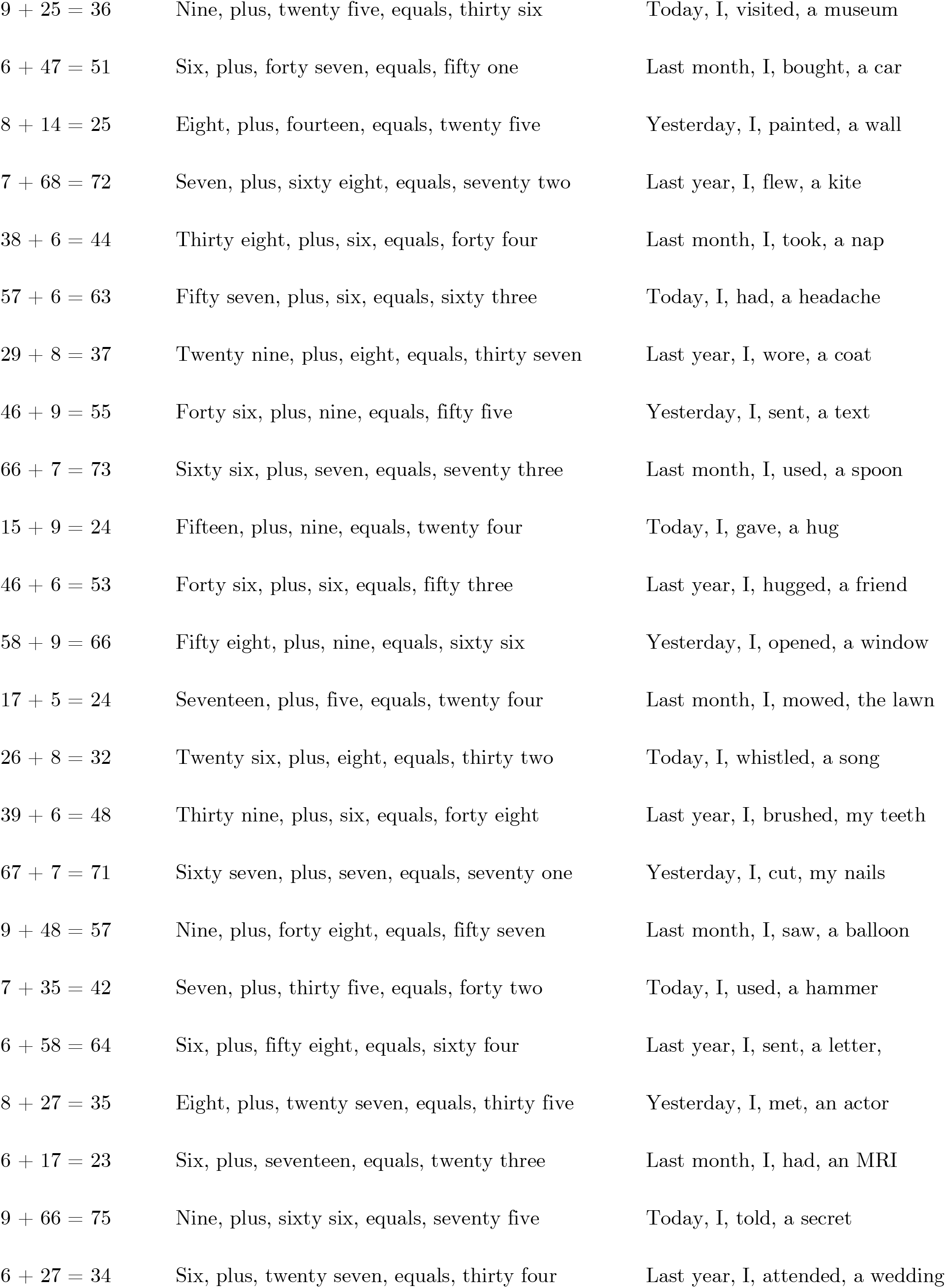
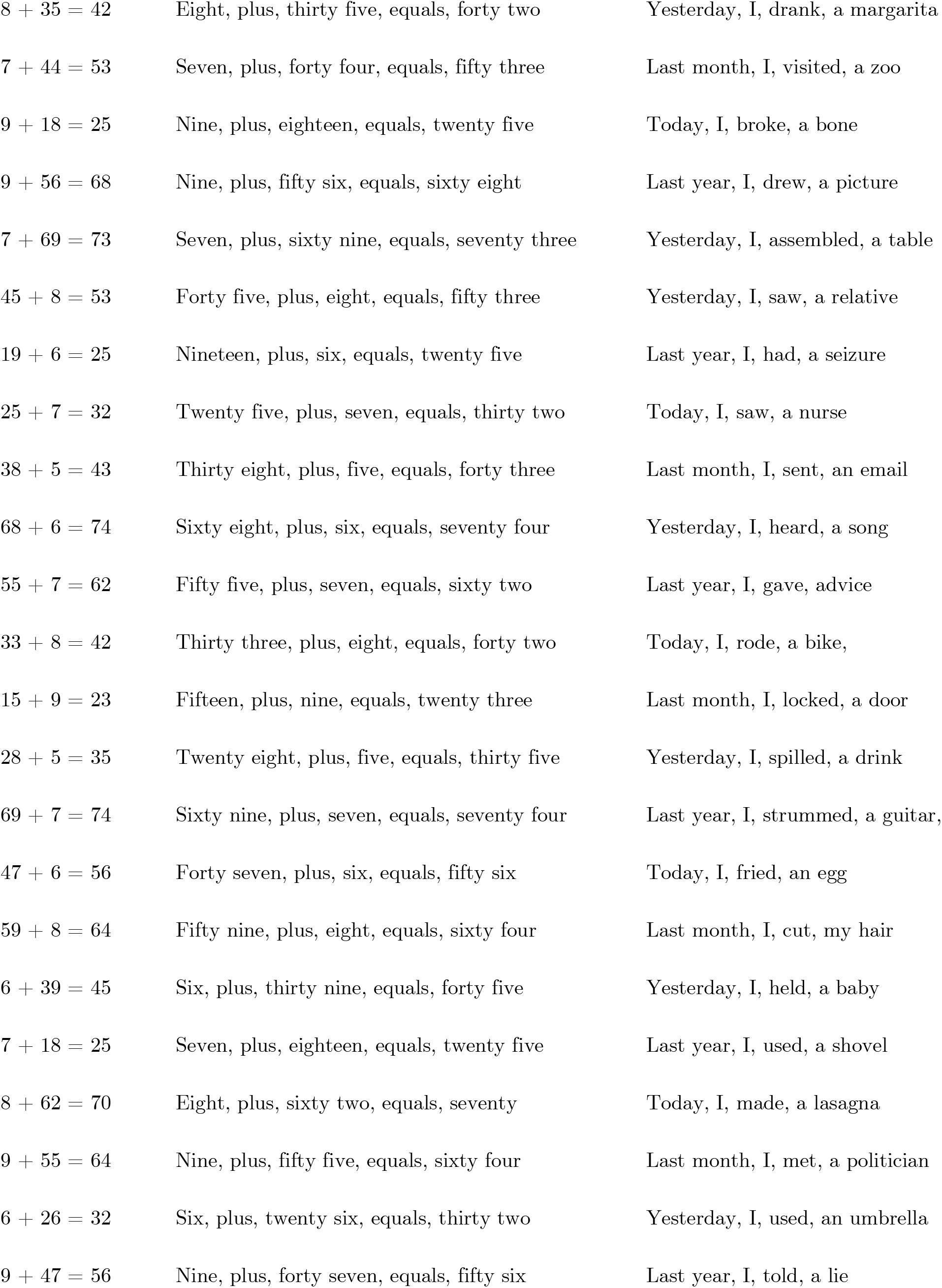
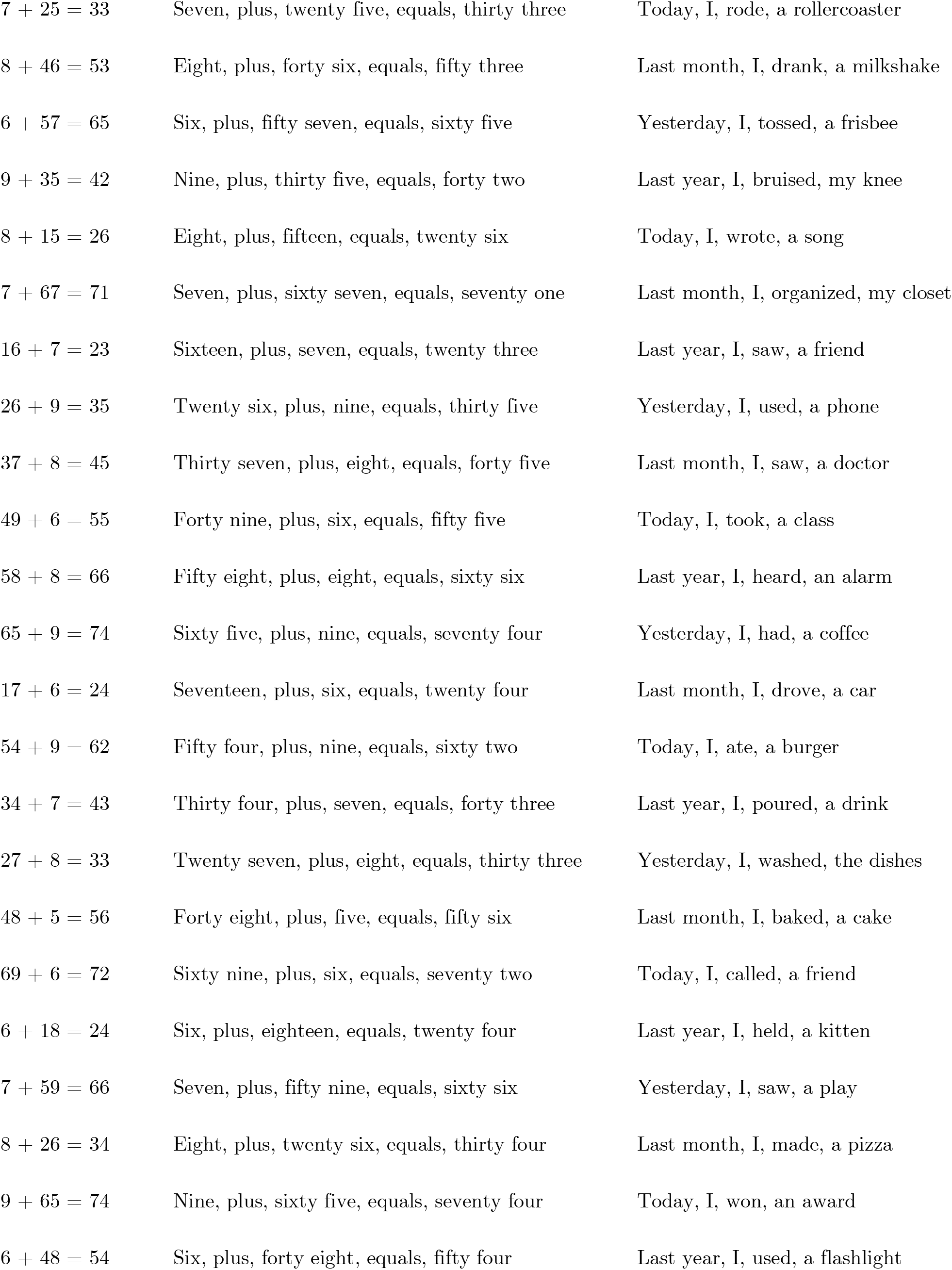
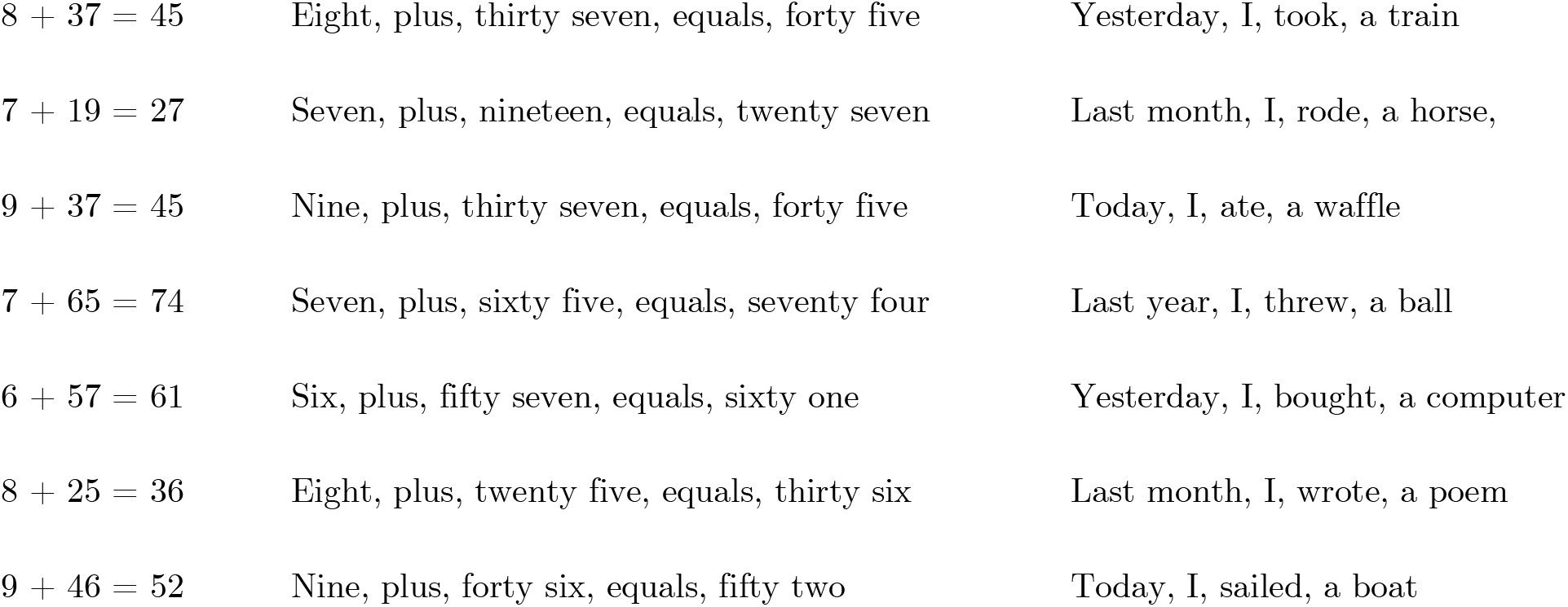
Stimuli list for the Sequential Calculation task.

**Figure 1-4.**
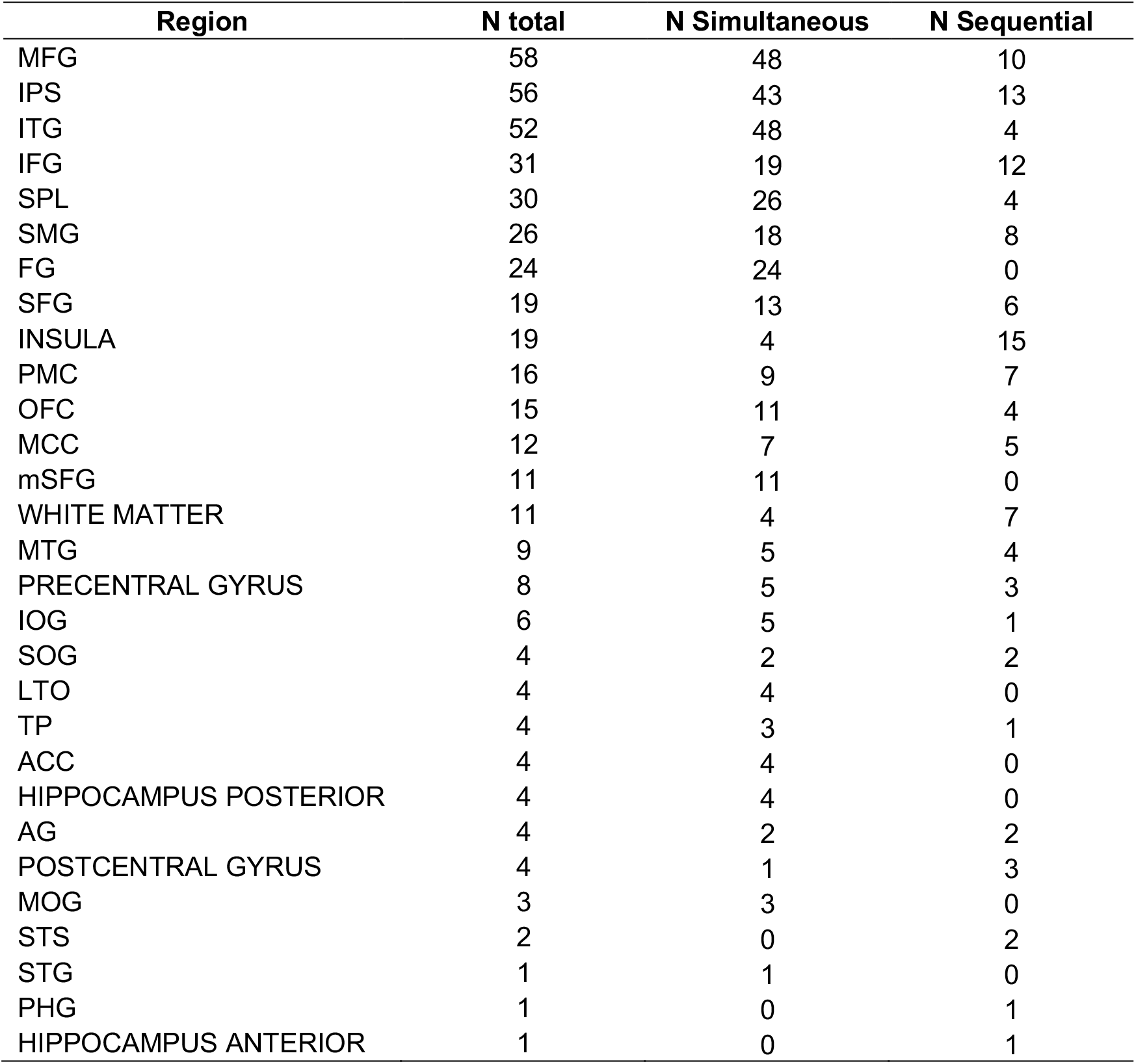
Math preferential sites per anatomical region.

**Figure 1-5.**
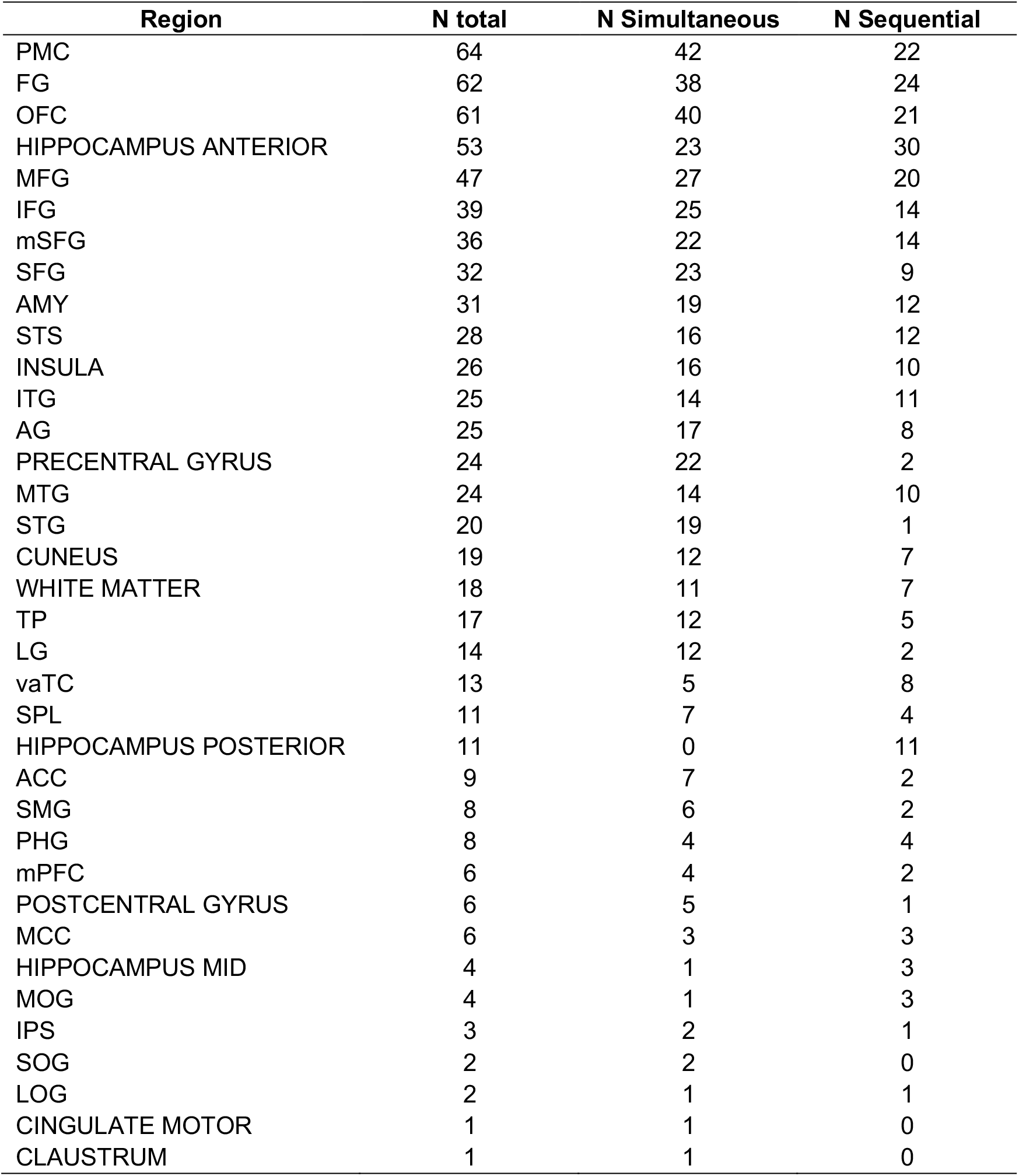
Non-math preferential sites per anatomical region.

**Figure 1-6.**
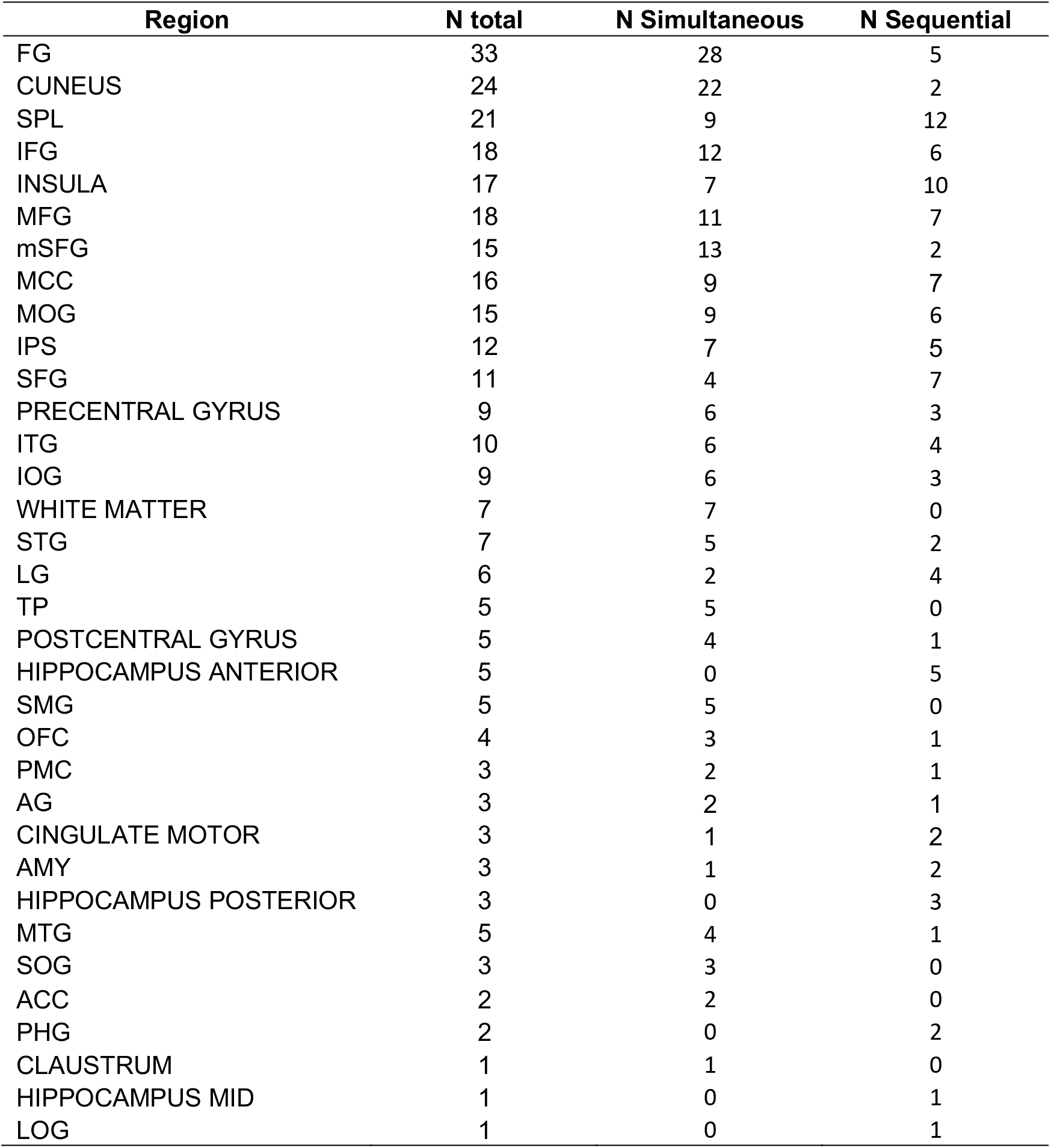
Math and Non-math equally preferential sites per anatomical region.

**Figure 1-7.**
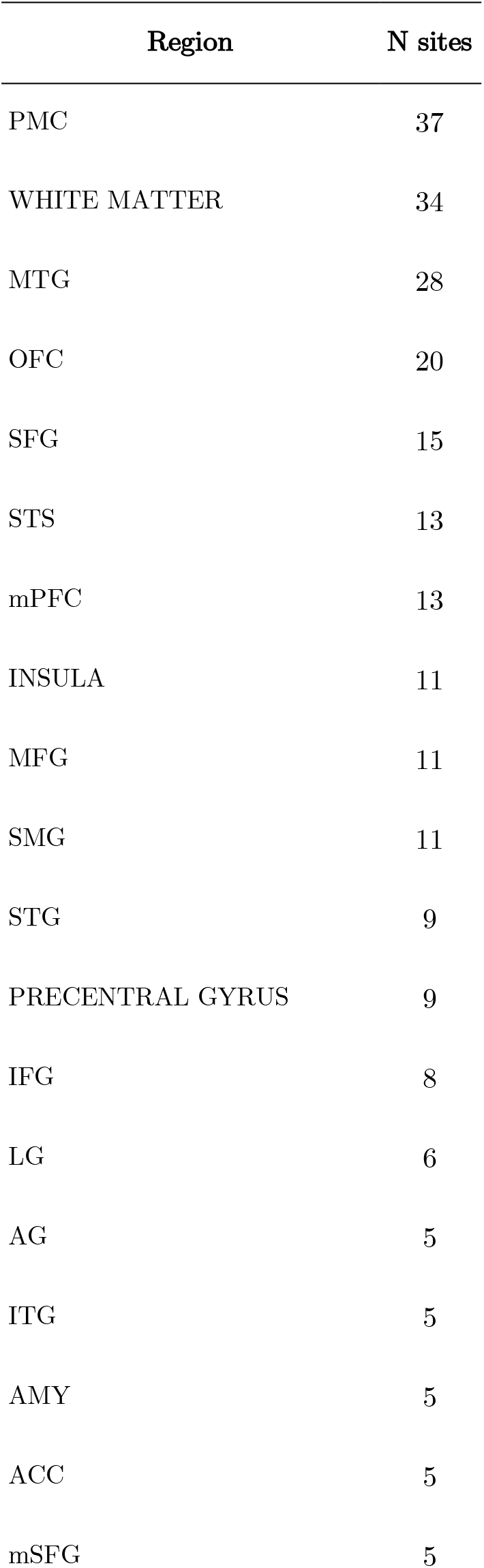
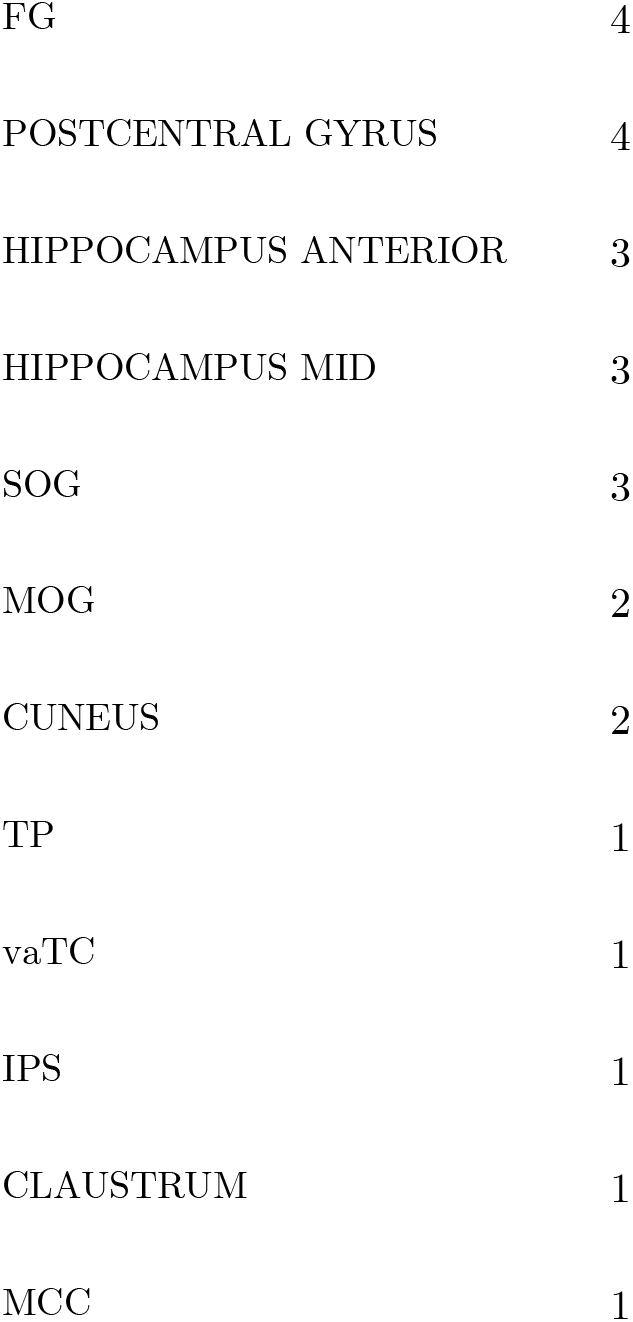
Math deactivated sites per anatomical region combining the Simultaneous and Sequential calculation tasks results.

**Figure 1-8.**
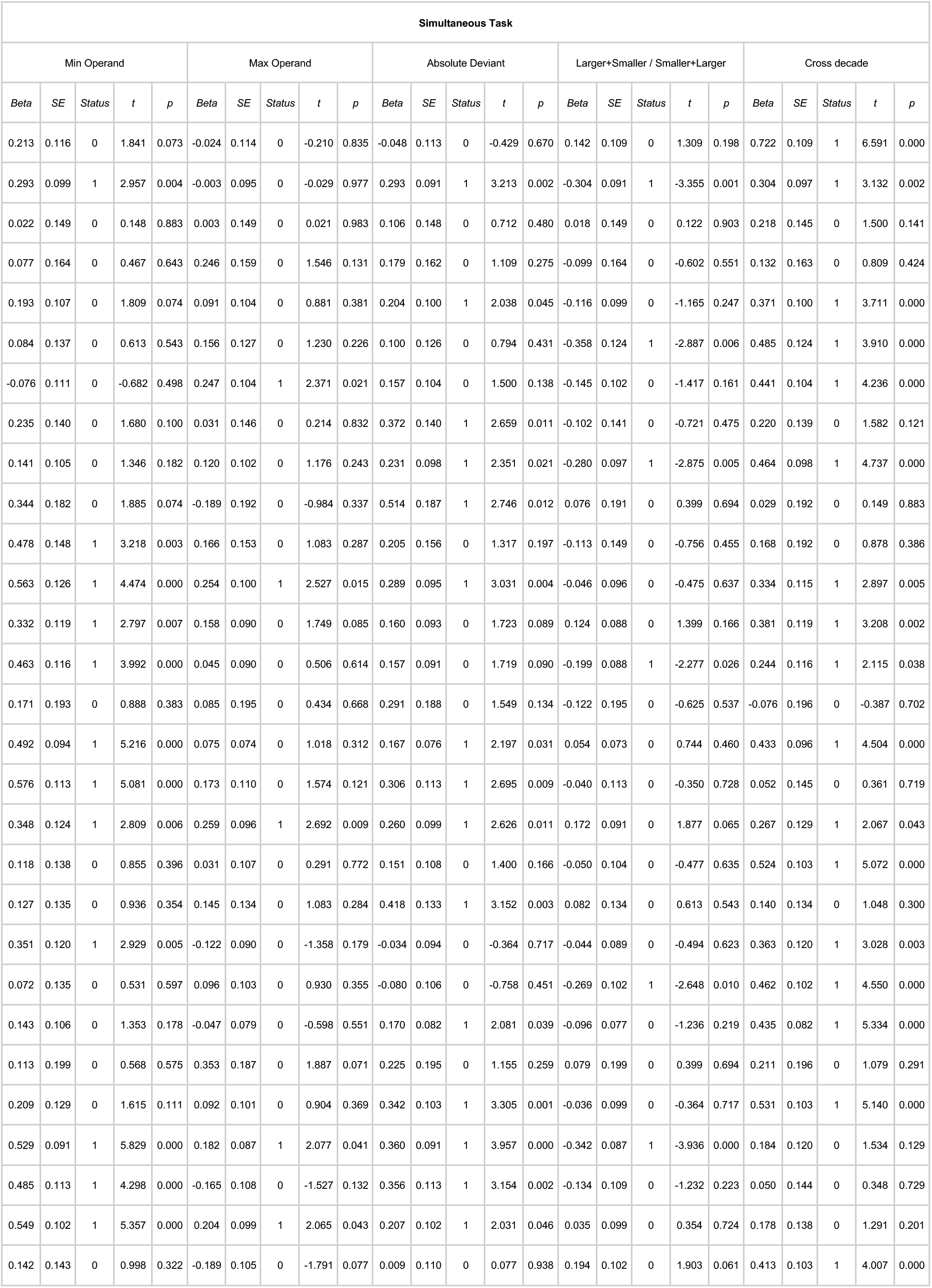
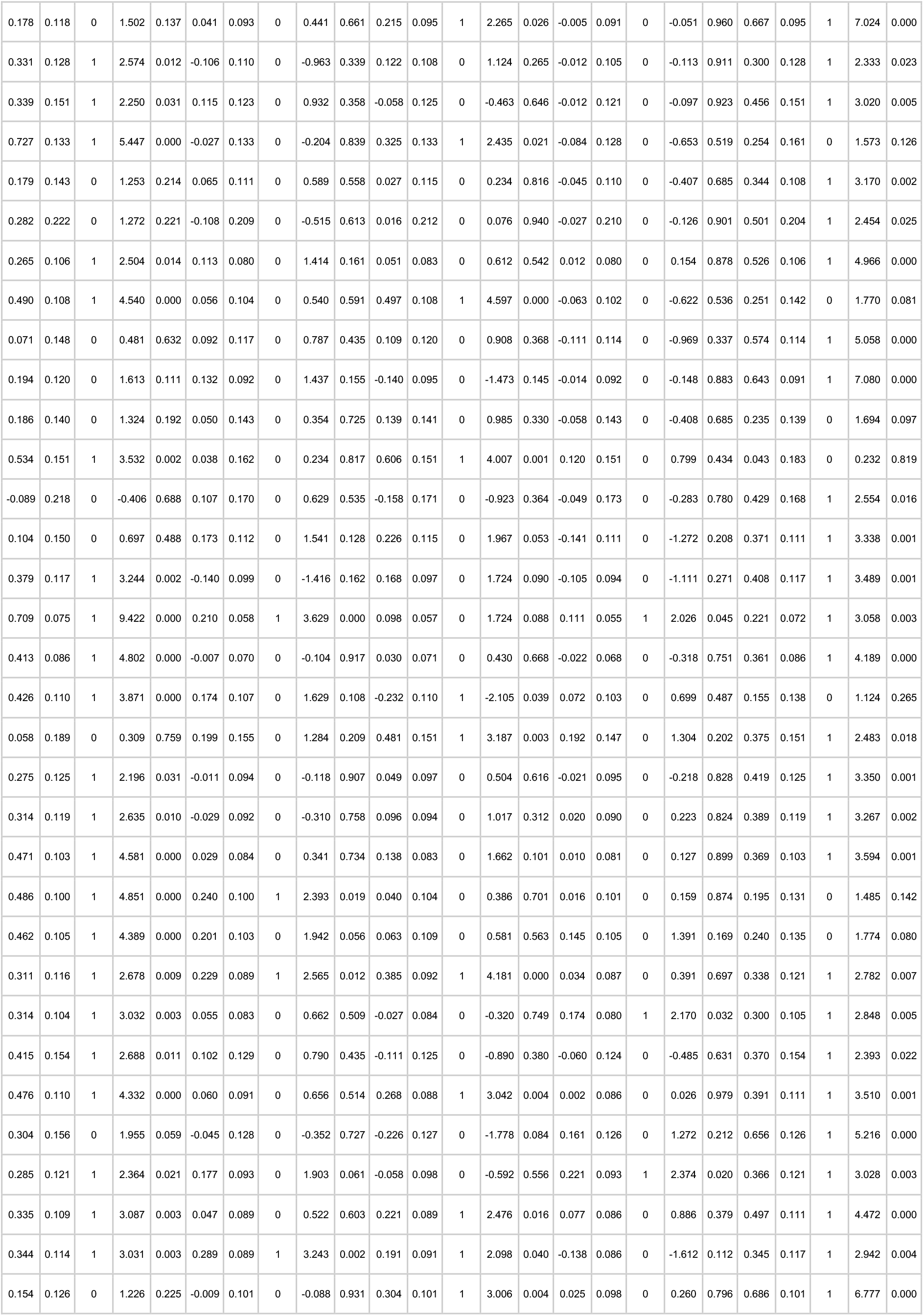
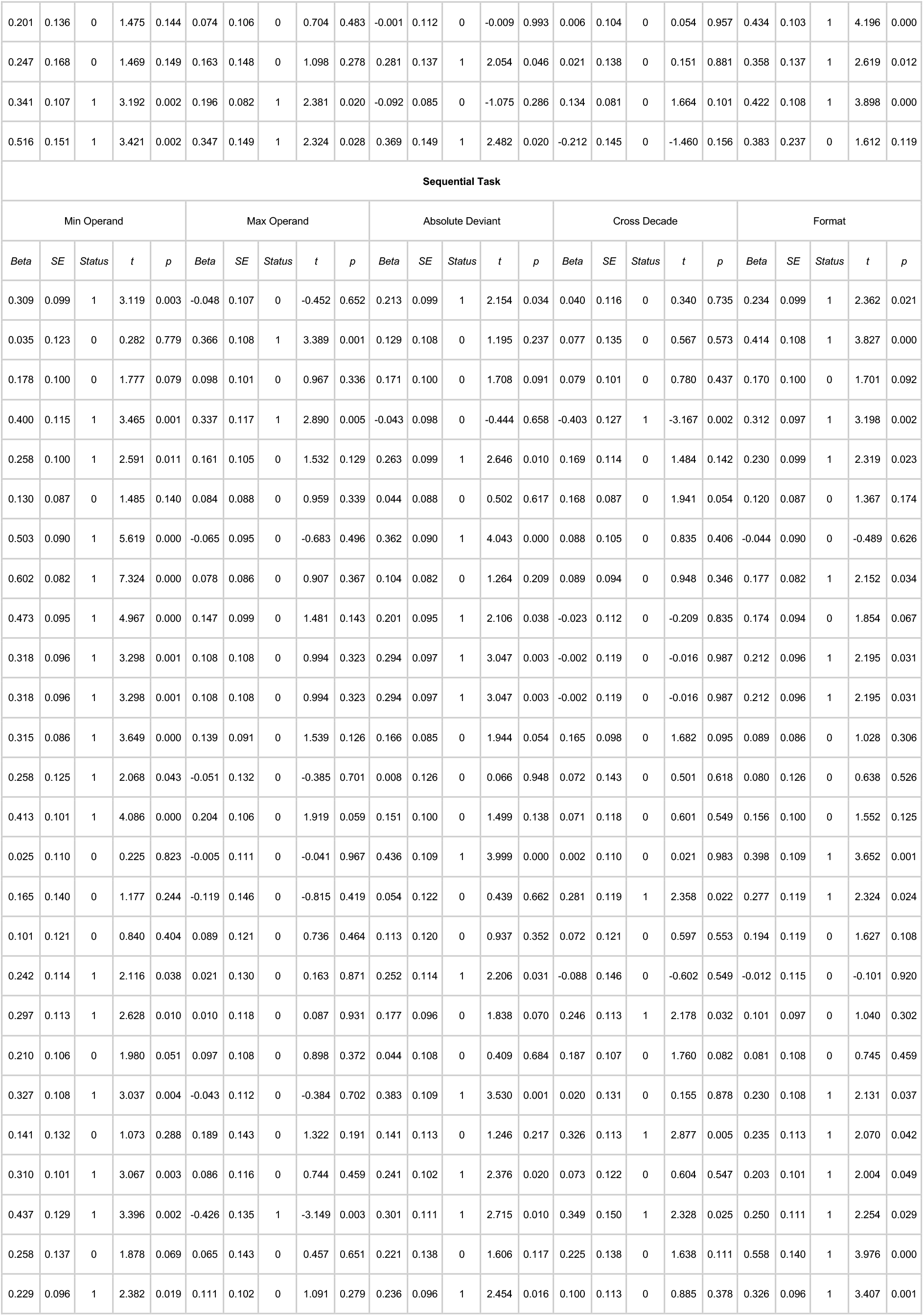
Arithmetic problem-size effect by subject. Supports Figure 1.

